# Relative demographic susceptibility does not explain the extinction chronology of Sahul’s megafauna

**DOI:** 10.1101/2020.10.16.342303

**Authors:** Corey J. A. Bradshaw, Christopher N. Johnson, John Llewelyn, Vera Weisbecker, Giovanni Strona, Frédérik Saltré

## Abstract

The causes of Sahul’s megafauna extinctions remain uncertain, although multiple, interacting factors were likely responsible. To test hypotheses regarding plausible ecological mechanisms underlying these extinctions, we constructed the first stochastic, age-structured models for 13 extinct megafauna species from five functional/taxonomic groups, as well as 8 extant species within these groups for comparison. Perturbing specific demographic rates individually, we tested which species were more demographically susceptible to extinction, and then compared these relative sensitivities to the fossil-derived extinction chronology. Here we show that the macropodiformes were the most resilient to extinction, followed by carnivores, monotremes, vombatiform herbivores, and large birds. Five of the eight extant species were as or more susceptible than were the extinct species. There was no clear relationship between extinction susceptibility and the extinction chronology for any perturbation scenario, but body mass and generation length explained much of the variation in relative risk. Our models reveal that the actual mechanisms leading to extinction were unlikely related to variation in demographic susceptibility *per se*, but were driven instead by finer-scale variation in climate change and/or human prey choice and relative hunting success.

## Introduction

The myriad mechanisms driving species extinctions [1] are often synergistic [2], spatially variable [3], phylogenetically clumped [4], correlated with population size [5], and dependent on biotic interactions [6]. This complexity means that even in contemporary settings involving closely monitored species, identifying the ecological mechanisms underlying the causes of a particular extinction can be difficult [7, 8]. This challenge is considerably greater for palaeo-extinctions because of the restricted ecological knowledge about extinct species. In such cases, we can only infer the conditions likely operating at the estimated time of disappearance from rare and sparsely distributed proxies.

The rapid and widespread disappearance of megafauna in the late Quaternary on most continents is one of the best-studied mass-extinction events of the past, and had many plausible causes [9]. The main drivers of extinction appear to differ in each case depending on taxa, region and time period [10–12], but there is growing consensus that multiple drivers were involved, including the interaction between climatic shifts and novel human pressure as dominant mechanisms [3, 13–16]. This consensus mostly relies on approaches examining extinction chronologies relative to indices of temporal and spatial environmental variation. While such correlative approaches can suggest potential causes of extinction, they cannot by themselves provide strong inference on the plausible ecological processes involved. Instead, approaches that construct mechanistic models of environmental and other processes that drive extinctions could hypothetically reveal the relative susceptibility of species over the course of a large extinction event [17].

Existing mechanistic models applied to megafauna systems differ in their complexity, ranging from predator-prey models [18, 19], to fully age-structured stochastic models [20], or stochastic predator-prey-competition functions [21]. If sufficiently comprehensive, such models can be useful tools to test hypotheses about the processes of extinction virtually in long-disappeared species. Although measuring the demographic rates of long-extinct species is impossible, robust rates can be inferred approximately from modern analogues and allometric relationships derived from extant species [20, 21]. Therefore, it is possible to construct stochastic demographic models of both extinct and related extant species, and compare their relative susceptibility to perturbations by mimicking particular environmental processes *in silico*.

Despite these examples of methodological advance, unravelling the causes underlying the disappearance of megafauna from Sahul (the combined landmass of Australia and New Guinea joined during periods of low sea level) is still a major challenge given the event’s antiquity [16] and the sparse palaeo-ecological information available relative to megafauna extinctions nearly everywhere else in the world [3, 16]. However, based on the expectation that if high demographic susceptibility is an important feature of a species’ actual extinction dynamics, the most susceptible species should have gone extinct before more resilient species did.

Stochastic demographic models can therefore potentially test the relative contribution of the following five mechanisms regarding the putative drivers of the megafauna extinctions in Sahul (summarized in Fig. 1): (***i***) There is a life history pattern in which the slowest-reproducing species succumbed first to novel and efficient human hunting [1, 22, 23]. This hypothesis assumes that human hunting, even if non-selective, would differentially remove species that were more demographically sensitive to increased mortality arising from novel human exploitation [24]. (***ii***) The most susceptible species were those whose life habits conferred the highest exposure to human hunters, such as species with vulnerable juveniles [25], or those occupying semi-open habitats like savannas, compared to those living in denser forests or in more inhospitable terrain (e.g., swamplands, mountains) less accessible to human hunters [24]. (***iii***) Bottom-up processes drove the extinctions, manifested as a difference in the timing of extinction between carnivores and their herbivore prey. Under this mechanism, prey-specialist carnivores should be more susceptible than their prey (i.e., because they depend on declining prey populations), whereas more generalist carnivores that can switch food sources would be less susceptible than their main prey [26, 27]. (***iv***) Species susceptible to temporal variation in climate would succumb before those most able to adapt to changing conditions. Under this hypothesis, we expect the largest species — i.e., those possessing traits associated with diet/habitat generalism [28], physiological resilience to fluctuating food availability [29, 30], high fasting endurance, and rapid, efficient dispersal away from stressful conditions [31] — would persist the longest in the face of catastrophic environmental change, independent of the intensity of human predation. (***v***) If none of the aforementioned mechanisms explains the extinction event’s chronology, non-demographic mechanisms such as differential selection of or ease of access by human hunters could have played more important roles.

**Figure 1.**
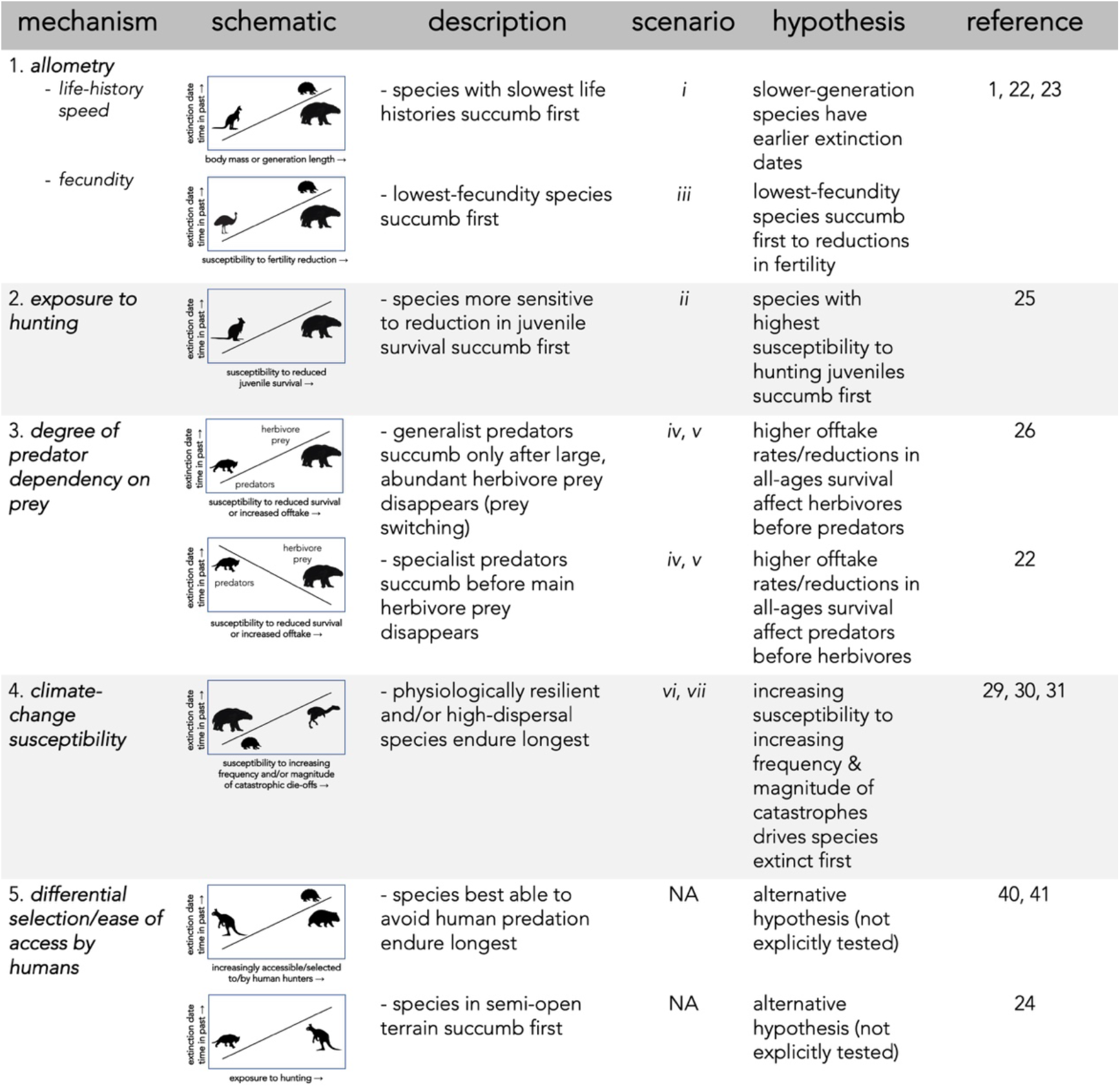
Description of five dominant mechanisms by which megafauna could have been driven to extinction and the associated hypotheses tested in Scenarios 1–7.

To test these various hypotheses, we developed the first stochastic, age-structured demographic models ever constructed for thirteen extinct megafauna species in Sahul broadly categorized into five functional/taxonomic groups: (*i*) four vombatiform herbivores, (*ii*) five macropodiform herbivores, (*iii*) one large, flightless bird, (*iv*) two marsupial carnivores, and (*v*) one monotreme invertivore. We also built demographic models for eight of some of the largest extant species, including representatives from each of the functional/taxonomic groups described above for comparison. Our null hypothesis is that these extant species should demonstrate higher resilience to perturbations than the extinct species, given that they persisted through the main extinction event to the present. Subjecting each species’ model stochastically to different scenarios of demographic perturbations, we tested seven scenarios (described in more details in *Methods*) regarding the processes that could lead to extinction (see also Fig. 1): (*i*) an allometric relationship between the time of extinction and species’ body mass and/or generation length, (*ii*) increasing juvenile mortality [25, 32], (*iii*) reducing fertility [33–35], (*iv*) reducing survival across all ages, (*v*) animal offtake from the population via hunting, (*vi*) increasing environmental variability generating extreme climate events, and (*vii*) increasing environment-driven catastrophic mortality events [36].

We hypothesize that one, or several, of these types of perturbations would provide a better match between relative demographic susceptibility and the continental-scale chronology of extinctions as inferred from the fossil record. Identifying which, if any, of the scenarios best matches the chronology would therefore indicate higher relative support for those mechanisms being the most likely involved in driving the observed extinctions. We first compared the expectation of larger (and therefore, slower life-history; Scenario *i*, Fig. 1) species more likely to go extinct than smaller species when faced with novel mortality sources [37], followed by the outcomes of all other scenarios (Scenarios *ii–vii*, Fig. 1) to test if sensitivity to specific demographic changes supported other mechanisms.

## Results

There was no indication that relatively heavier (Fig. 2a) or slower life history (longer-generation; Fig. 2b) species went extinct before lighter, faster life-history species (Scenario *i*), even considering that the two mid- and small-sized carnivores *Thylacinus* and *Sarcophilus* went extinct on the mainland late in the Holocene at approximately the same time (~ 3200 years before present; Fig. 2) [38].

**Figure 2.**
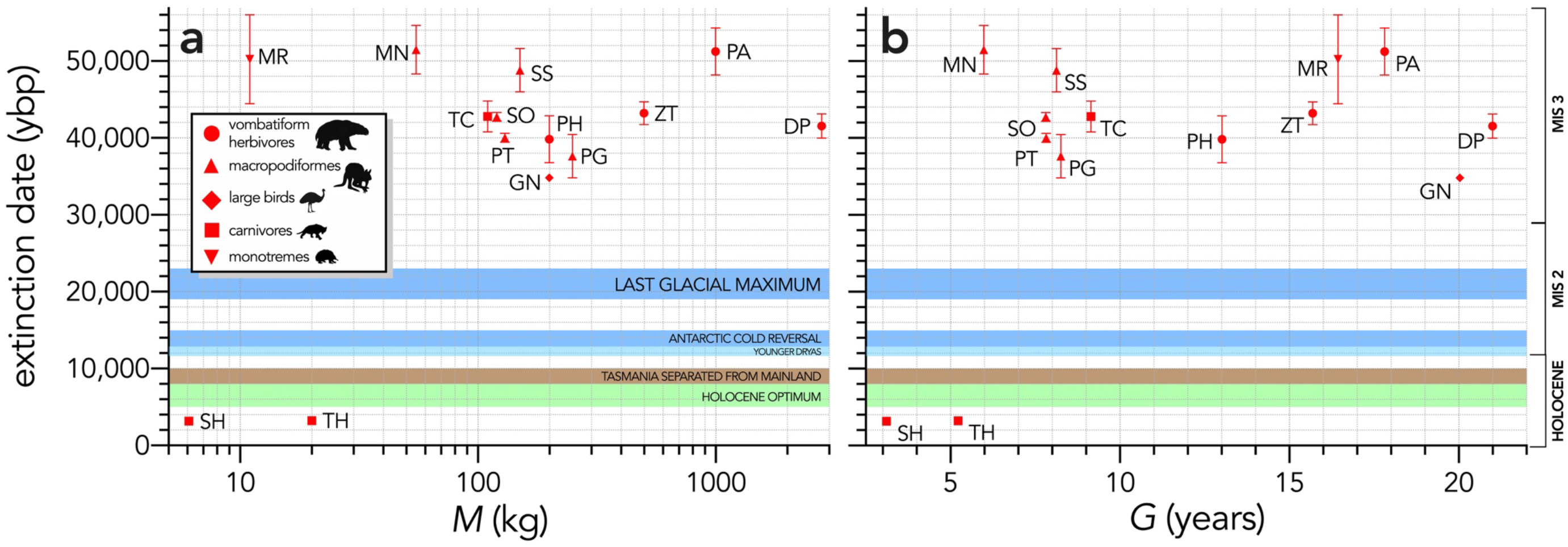
Relationship between estimated date of species extinction (across entire continent) and (**a**) body mass (kg) or (**b**) generation length (years) (Scenario 1). Species notation: DP = *Diprotodon optatum*, PA = *Palorchestes azael*, ZT = *Zygomaturus trilobus*, PH = *Phascolonus gigas*, VU *Vombatus ursinus* (**vombatiform herbivores**); PG = *Procoptodon goliah*, SS = *Sthenurus stirlingi*, PT = *Protemnodon anak*, SO = *Simosthenurus occidentalis*, MN = *Metasthenurus newtonae*, OR = *Osphranter rufus* (**macropodiformes**); GN = *Genyornis newtoni*, DN = *Dromaius novaehollandiae* (**large birds**); TC = *Thylacoleo carnifex*, TH = *Thylacinus cynocephalus*, SH = *Sarcophilus harrisii* (**carnivores**); TA = *Tachyglossus aculeatus*, MR = *Megalibgwilia ramsayi* (**monotreme invertivores**). Here, we have depicted SH as ‘extant’, even though it went extinct on the mainland > 3000 years ago. Also shown are the approximate major climate periods and transitions: Marine Isotope Stage 3 (MIS 3), MIS 2 (including the Last Glacial Maximum, Antarctic Cold Reversal, and Younger Dryas), and the Holocene (including the period of sea level flooding when Tasmania separated from the mainland, and the relatively warm, wet, and climatically stable Holocene optimum).

The quasi-extinction curves (used as a proxy for extinction risk) for each species differed markedly in each perturbation scenario (Supplementary Information Appendix S6, Fig. S7), although there were some similarities among scenarios. For example, in all scenarios except for fertility-reduction (Scenario *iii*) and offtake (Scenario *v*), the smallest extant carnivore *Dasyurus* was the least susceptible, whereas *Genyornis* was one of the most susceptible in 4 of the 6 scenarios (Supplementary Information Appendix 5, Fig. S7).

Across all species, log_10_ body mass explained some of the variance in the total area under the quasi-extinction curve (Fig. 3) for individual removal (evidence ratio [ER] = 41.22, *R^2^* = 0.35; Scenario *v*; Fig. 3d) and catastrophe magnitude (ER = 19.92, *R*^2^ = 0.31; Scenario *vii*; Fig. 3f), but less variance for reductions in juvenile (ER = 2.04, *R*^2^ = 0.14; Scenario *ii*; Fig. 3a) and all-ages survival (ER = 5.05, *R*^2^ = 0.21; Scenario *iv*; Fig. 3c). There was little to no evidence for a relationship in the fertility-reduction and catastrophe-frequency scenarios (ER ≲ 1; Fig. 3b, e). The relationships were generally stronger between area under the quasi-extinction curve and log_10_ generation length (*G*) (Fig. 4). The strongest relationships here were for all-ages survival reduction and magnitude of catastrophe (ER > 490, *R*^2^ ≥ 0.49; Scenarios *iv* and *vii*; Fig. 4c, f), followed by weaker relationships (ER < 11, *R*^2^ ≤ 0.26) for Scenarios *ii* (Fig. 4a), *v* (Fig. 4d), and *vi* (Fig. 4e), and no evidence for a relationship in Scenario *iii* (ER < 1; Fig. 4b).

**Figure 3.**
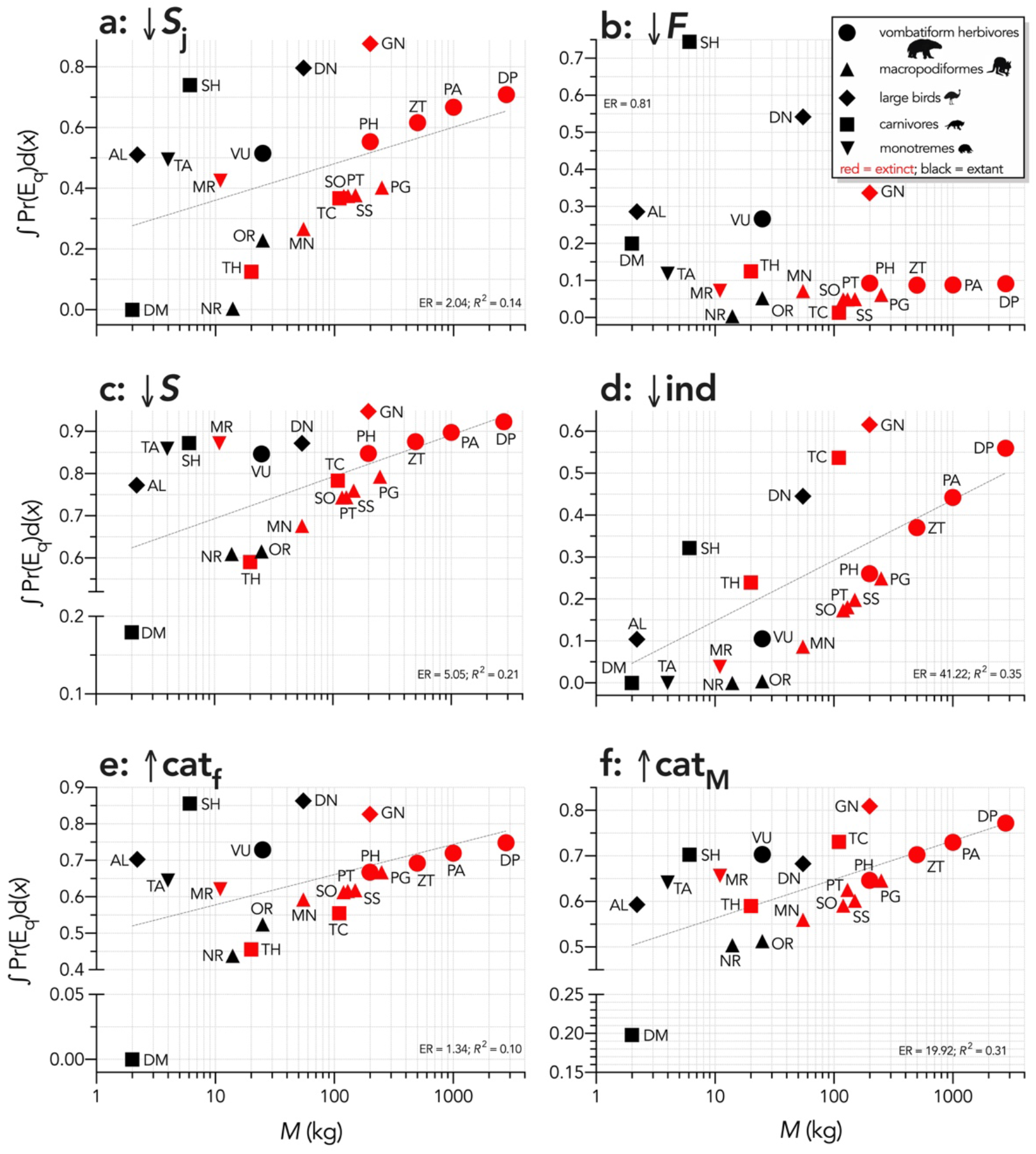
Area under the quasi-extinction curve (from Fig. S7) — ∫Pr(E_q_)d(*x*) — as a function of body mass (*M*, kg) for **a**: (↓***S*_j_**) increasing juvenile mortality (Scenario *ii*), **b**: (↓***F***) decreasing fertility (Scenario *iii*), **c**: (↓***S***) decreasing survival across all age classes (Scenario *iv*), **d**: (↓**ind**) increasing number of individuals removed year^-1^ (Scenario *v*), **e**: (↑**cat_f_**) increasing frequency of catastrophic die-offs per generation (Scenario *vi*), and **f**: (↑**cat_M_**) increasing magnitude of catastrophic die-offs (Scenario *vii*). Shown are the information-theoretic evidence ratios (ER) and variation explained (*R^2^*) for the lines of best fit (grey dashed) in each scenario. Species notation: DP = *Diprotodon optatum*, PA = *Palorchestes azael*, ZT = *Zygomaturus trilobus*, PH = *Phascolonus gigas*, VU *Vombatus ursinus* (**vombatiform herbivores**); PG = *Procoptodon goliah*, SS = *Sthenurus stirlingi*, PT = *Protemnodon anak*, SO = *Simosthenurus occidentalis*, MN = *Metasthenurus newtonae*, OR = *Osphranter rufus*, NR = *Notamacropus rufogriseus*, common name: red-necked wallaby (**macropodiformes**); GN = *Genyornis newtoni*, DN = *Dromaius novaehollandiae*, AL = *Alectura lathami* (**large birds**); TC = *Thylacoleo carnifex*, TH = *Thylacinus cynocephalus*, SH = *Sarcophilus harrisii*, DM = *Dasyurus maculatus* (**carnivores**); TA = *Tachyglossus aculeatus*, MR = *Megalibgwilia ramsayi* (**monotreme invertivores**). Here, we have depicted SH as ‘extant’, even though it went extinct on the mainland > 3000 years ago.

**Figure 4.**
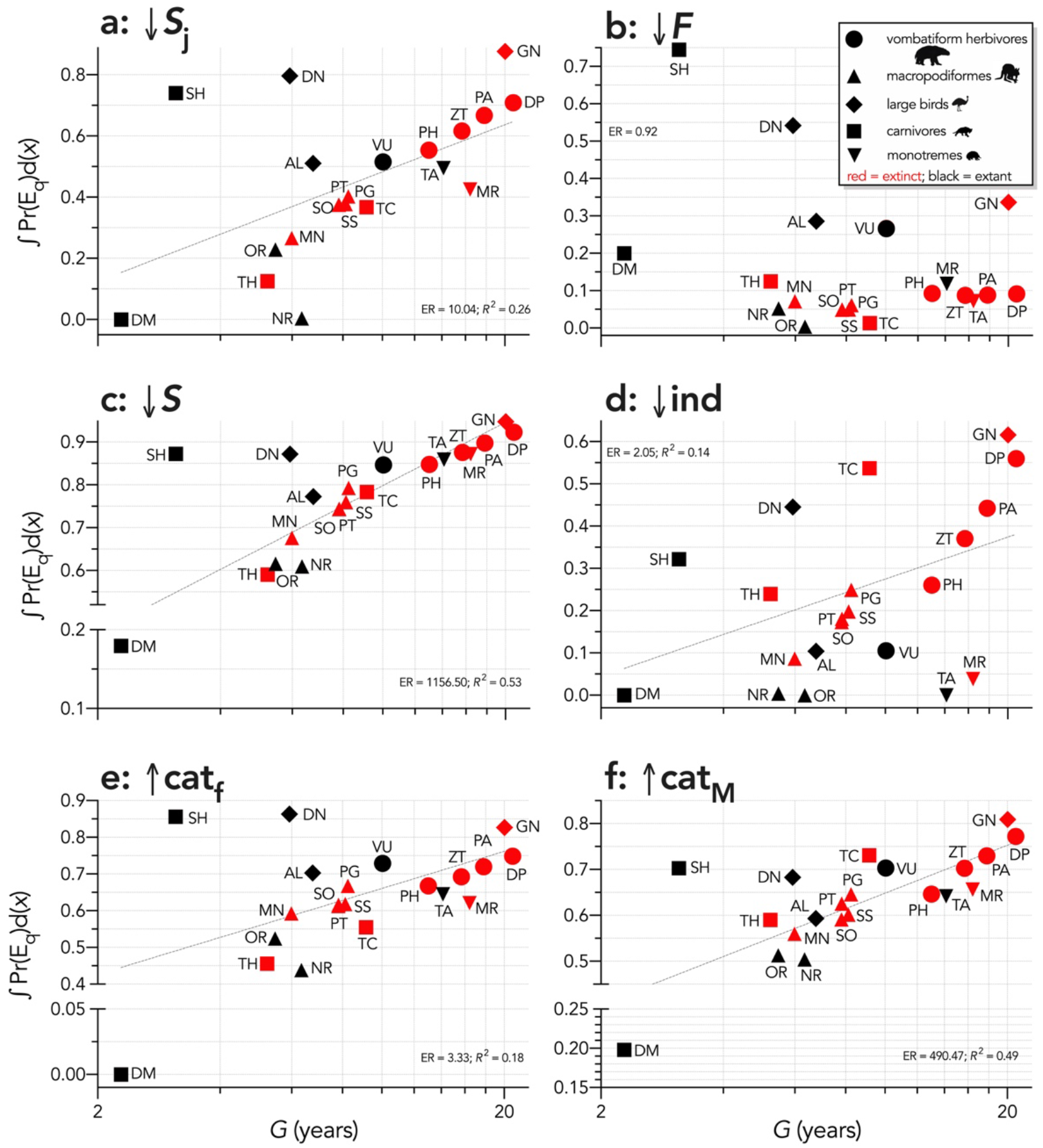
Area under the quasi-extinction curve (from Fig. S7) — ∫Pr(E_q_)d(*x*) — as a function of generation length (*G*, years) for **a**: (↓***S***_**j**_) increasing juvenile mortality (Scenario *ii*), **b**: (↓***F***) decreasing fertility (Scenario *iii*), **c**: (↓***S***) decreasing survival across all age classes (Scenario *iv*), **d**: (↓**ind**) increasing number of individuals removed year^-1^ (Scenario *v*), **e**: (↑**cat_f_**) increasing frequency of catastrophic die-offs per generation (Scenario *vi*), and **f**: (↑**cat_M_**) increasing magnitude of catastrophic die-offs (Scenario 7). Shown are the information-theoretic evidence ratios (ER) and variation explained (*R^2^*) for the lines of best fit (grey dashed) in each scenario. Species notation: DP = *Diprotodon optatum*, PA = *Palorchestes azael*, ZT = *Zygomaturus trilobus*, PH = *Phascolonus gigas*, VU *Vombatus ursinus* (**vombatiform herbivores**); PG = *Procoptodon goliah*, SS = *Sthenurus stirlingi*, PT = *Protemnodon anak*, SO = *Simosthenurus occidentalis*, MN = *Metasthenurus newtonae*, OR = *Osphranter rufus*, NR = *Notamacropus rufogriseus*, common name: red-necked wallaby (**macropodiformes**); GN = *Genyornis newtoni*, DN = *Dromaius novaehollandiae*, AL = *Alectura lathami* (**large birds**); TC = *Thylacoleo carnifex*, TH = *Thylacinus cynocephalus*, SH = *Sarcophilus harrisii*, DM = *Dasyurus maculatus* (**carnivores**); TA = *Tachyglossus aculeatus*, MR = *Megalibgwilia ramsayi* (**monotreme invertivores**). Here, we have depicted SH as ‘extant’, even though it went extinct on the mainland > 3000 years ago.

Allometric scaling of risk was also apparent within most taxonomic/functional groups. For example, *Diprotodon* had the highest extinction risk among the extinct vombatiform herbivores in every scenario except fertility reduction (Scenario *iii*; Fig. 3b, 4b). Most species were relatively immune to even large reductions in fertility, except for *Sarcophilus, Dasyurus, Vombatus, Alectura, Dromaius*, and *Genyornis* (Fig. 3b, 4b). Sub-Scenario *iib* where we removed only juvenile individuals was qualitatively similar to Scenario *ii* where we progressively increased juvenile mortality (Supplementary Information Appendix S7, Fig. S8), although the relative susceptibility for most species decreased from Scenario *ii* to *iib*. However, susceptibility increased for the extinct carnivores *Thylacinus* and *Thylacoleo*, and remained approximately the same for *Dasyurus* and *Notamacropus* (Fig. S8).

In the sub-Scenario *iiib* where we progressively increased the mean number of eggs removed per female per year in the bird species to emulate egg harvesting by humans, there was a progressively increasing susceptibility with body mass (Fig. 5). *Genyornis* was clearly the most susceptible to extinction compared to the other two bird species (Fig. 4).

**Figure 5.**
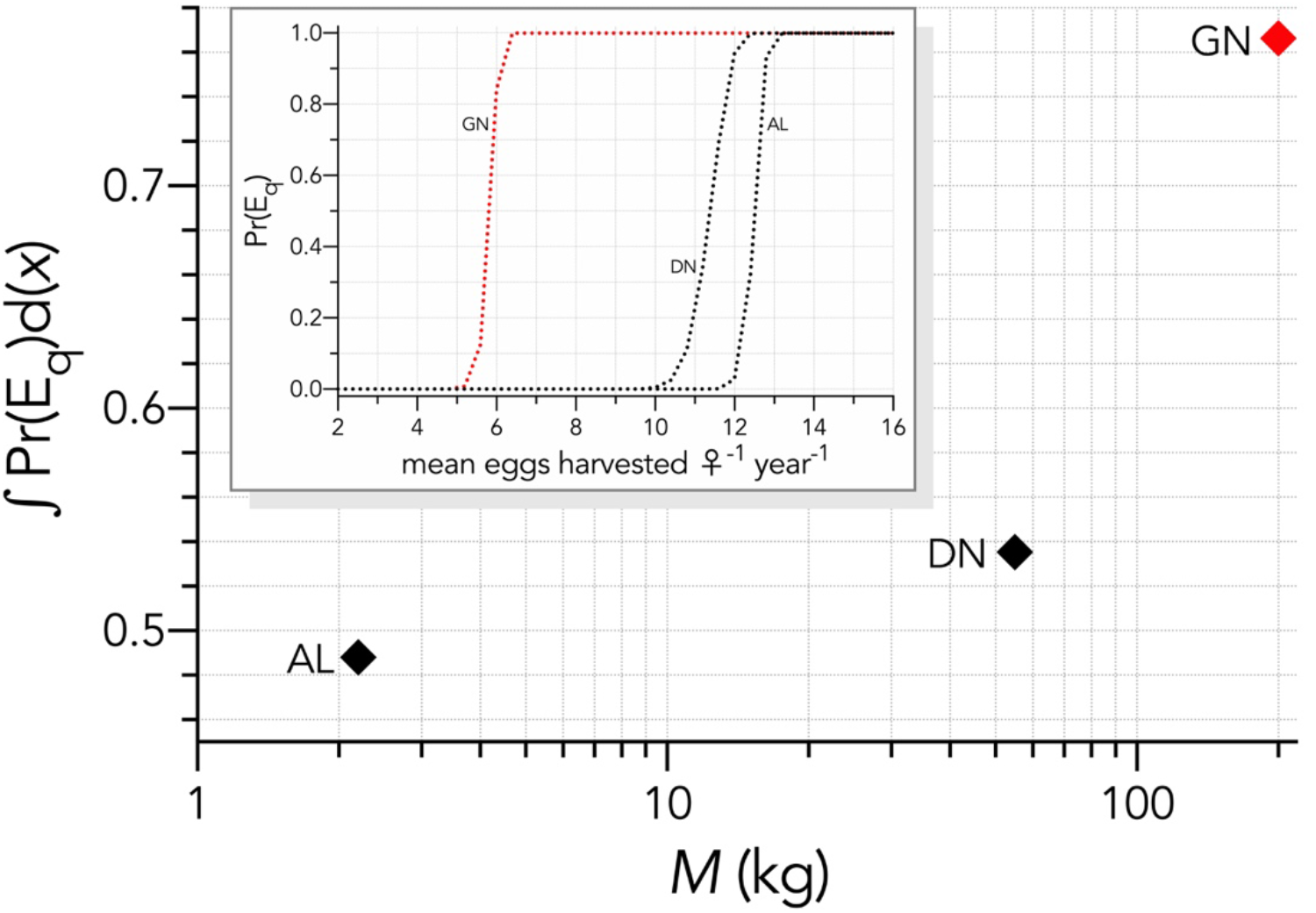
Inset: Increasing probabilities of quasi-extinction for birds — Pr(E_q_) — as a function of increasing the mean number of eggs harvested per female per year (Scenario *iiib*). The main graph shows the area under the quasi-extinction curve — ∫Pr(E_q_)d(*x*) — as a function body mass. Species notation: GN = *Genyornis newtoni*, DN = *Dromaius novaehollandiae*, AL = *Alectura lathami*.

When the extinction dates are viewed relative to the quasi-extinction integrals calculated for each scenario, there is no indication that the most susceptible species went extinct earlier in any perturbation scenario (Fig. 6). Taking the sum of the quasi-extinction integrals across scenarios indicated that five of the eight extant species examined (*Sarcophilus* [extinct on mainland; extant in Tasmania], *Dromaius, Alectura, Vombatus, Tachyglossus*) had extinction risks that were equivalent or higher compared to most of the extinct species (Fig. 7a). Taking the median rank of the quasi-extinction integral across scenarios generally indicated the highest resilience in the macropodiformes (although the small, extant carnivore *Dasyurus* was consistently top-ranked in terms of resilience for all scenarios except fertility reduction; Fig. 7a), followed by the carnivores (except *Dasyurus*), monotremes, vombatiform herbivores, and finally, large birds (Fig. 7b). The carnivores had resilience ranks spread across most of the susceptibility-rank spectrum (Fig. 7b).

**Figure 6.**
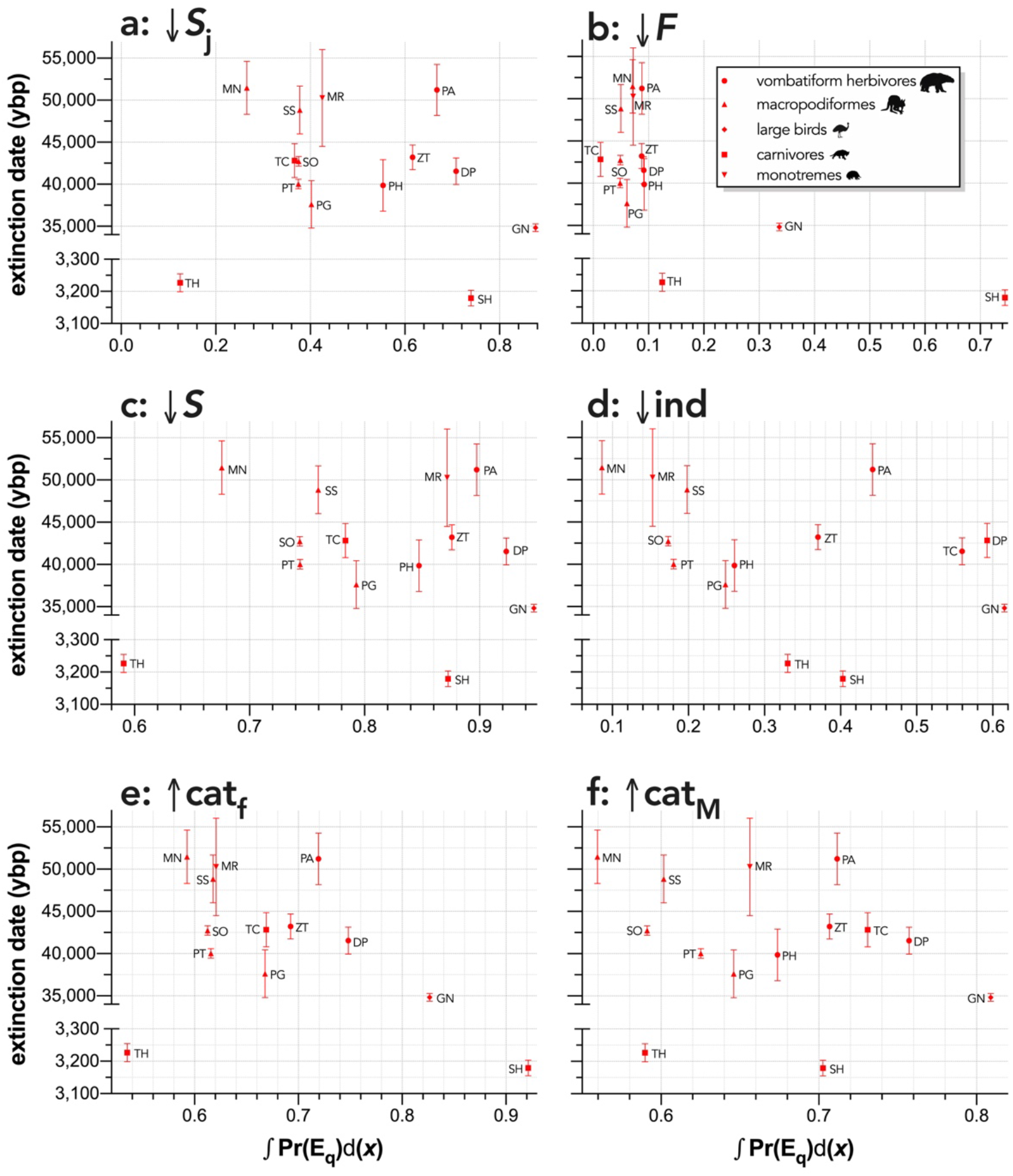
Relationship between estimated date of species extinction (across entire mainland) and area under the quasi-extinction curve (from Fig. S7) — ∫Pr(E_q_)d(*x*) — for **a**: (↓***S***_**j**_) increasing juvenile mortality (Scenario *ii*), **b**: (↓***F***) decreasing fertility (Scenario *iii*), **c**: (↓***S***) decreasing survival across all age classes (Scenario *iv*), **d**: (↓**ind**) increasing number of individuals removed year^-1^ (Scenario *v*), **e**: (↑**cat_f_**) increasing frequency of catastrophic die-offs per generation (Scenario *vi*), and **f**: (↑**cat_M_**) increasing magnitude of catastrophic die-offs (Scenario 7). Species notation: DP = *Diprotodon optatum*, PA = *Palorchestes azael*, ZT = *Zygomaturus trilobus*, PH = *Phascolonus gigas*, VU *Vombatus ursinus* (**vombatiform herbivores**); PG = *Procoptodon goliah*, SS = *Sthenurus stirlingi*, PT = *Protemnodon anak*, SO = *Simosthenurus occidentalis*, MN = *Metasthenurus newtonae*, OR = *Osphranter rufus*, NR = *Notamacropus rufogriseus*, common name: red-necked wallaby (**macropodiformes**); GN = *Genyornis newtoni*, DN = *Dromaius novaehollandiae*, AL = *Alectura lathami* (**large birds**); TC = *Thylacoleo carnifex*, TH = *Thylacinus cynocephalus*, SH = *Sarcophilus harrisii* (**carnivores**); TA = *Tachyglossus aculeatus*, MR = *Megalibgwilia ramsayi* (**monotreme invertivores**).

**Figure 7.**
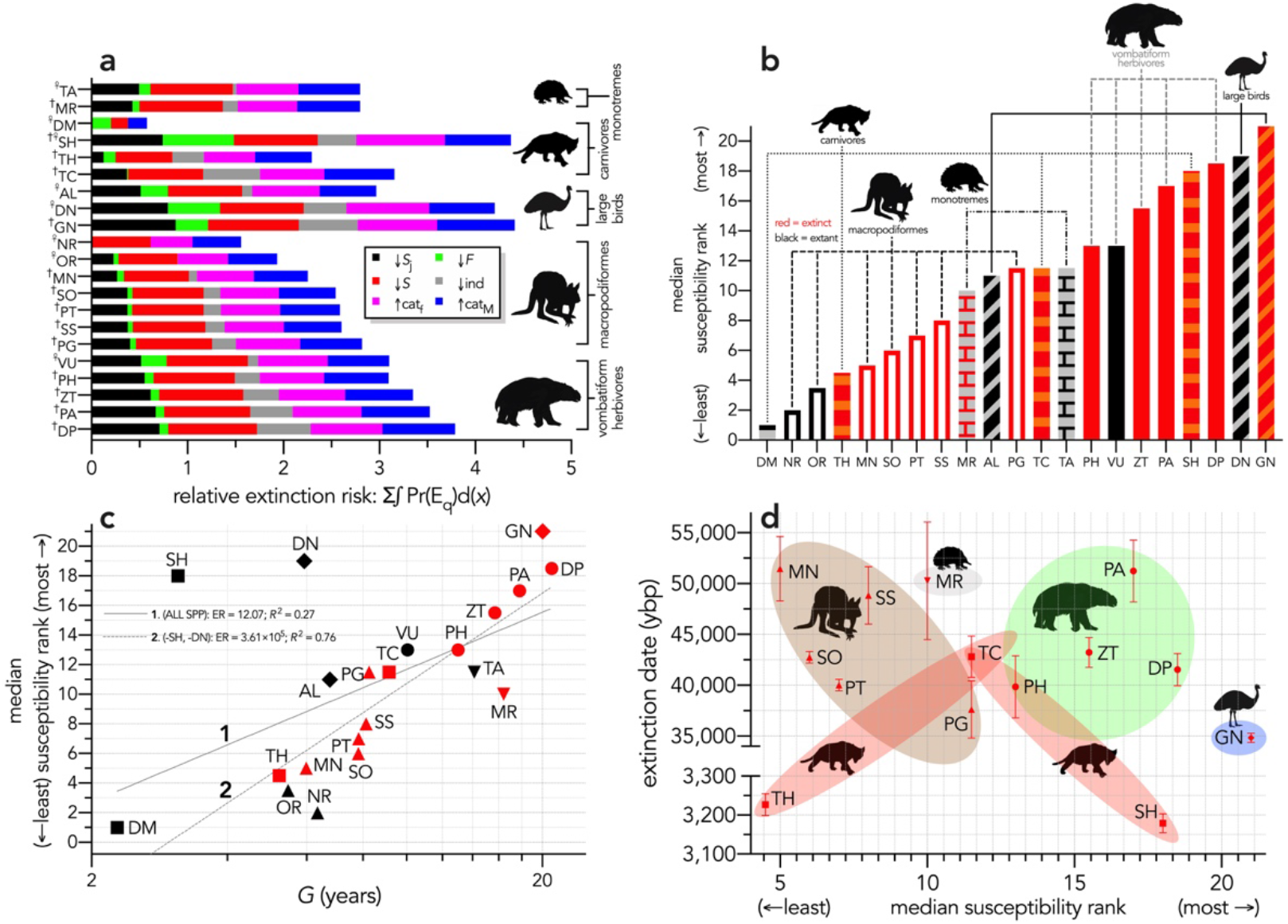
(**a**) Sum of the areas under the quasi-extinction curve for each of the six scenarios considered — Σ∫Pr(E_q_)d(*x*) — for each of the 21 modelled species (^†^extinct; ^♀^extant); (**b**) median resilience rank across the six scenarios considered (where lower ranks = higher resilience to extinction) for each species; (**c**) median resilience rank as a function of log_10_ generation length (*G*, kg) — there was a weak correlation including all species (solid grey line 1), but a strong relationship removing SH and DN (dashed grey line 2) (information-theoretic evidence ratio [ER] and variance explained [*R*^2^] shown for each); (**d**) estimated date of species extinction (across entire continent) as a function of median resilience rank; taxonomic/functional groupings are indicated by shaded colours (macropodids: brown; monotremes: grey; vombatiforms: green; birds: blue; carnivores: red). Species notation: DP = *Diprotodon optatum*, PA = *Palorchestes azael*, ZT = *Zygomaturus trilobus*, PH = *Phascolonus gigas*, VU *Vombatus ursinus* (**vombatiform herbivores**); PG = *Procoptodon goliah*, SS = *Sthenurus stirlingi*, PT = *Protemnodon anak*, SO = *Simosthenurus occidentalis*, MN = *Metasthenurus newtonae*, OR = *Osphranter rufus*, NR = *Notamacropus rufogriseus*, common name: red-necked wallaby (**macropodiformes**); GN = *Genyornis newtoni*, DN = *Dromaius novaehollandiae*, AL = *Alectura lathami* (**large birds**); TC = *Thylacoleo carnifex*, TH = *Thylacinus cynocephalus*, SH = *Sarcophilus harrisii*, DM = *Dasyurus maculatus* (**carnivores**); TA = *Tachyglossus aculeatus*, MR = *Megalibgwilia ramsayi* (**monotreme invertivores**). SH is extinct in mainland Australia, but extant in the island state of Tasmania.

Expressed as a function of log_10_ generation length, there was evidence for some relationship across all species (ER = 12.07, *R^2^* = 0.27), but removing the outlier species *Sarcophilus* and *Dromaius* resulted in a much stronger relationship (ER = 3.61×10^5^, *R*^2^ = 0.76). Susceptibility also tended to increase with body mass within a group, except for carnivores (*Sarcophilus* being the anomaly) and monotremes (Fig. 7c). There was no indication of a pattern when extinction date is plotted against median resilience rank (Fig. 7d); in fact, ignoring the late-surviving carnivores (*Thylacinus* and *Sarcophilus*) might suggest an opposite trend where the least demographically susceptible species went continentally extinct first in potential support of mechanism 4 tested in Scenarios *vi* and *vii* (Fig. 1), although the slope of this suggested relationship does not differ statistically from zero (randomization regression *p* = 0.272; see Supplementary Information Appendix S8 for a description of the randomized-regression method).

## Discussion

The megafauna species of Sahul demonstrate demographic susceptibility to extinction largely following expectations derived from threat risk in modern species — species with slower life histories have higher demographic risk to extinction on average [22]. Our twenty-one stochastic models indicate that the combined relative extinction risk of megafauna species in Sahul — taken in trophic isolation from the rest of the community — is not consistent with the estimated extinction chronology across continental Sahul. Actual extinction is instead an emergent property of many interacting demographic rates, temporal variation in population abundance, particular environmental contexts, and community interactions [1]. As different perturbations compromise different aspects of a species’ life history, its relative susceptibility to extinction compared to other species in its community varies in often unpredictable ways.

We can therefore reject the hypothesis that the continent-wide extinction chronology is explained by a species’ relative demographic susceptibility and non-selective hunting by humans. If demographic susceptibility coupled with non-selective hunting were the primary causes of these extinctions, we would expect a relationship between vulnerability and extinction date whereby the more vulnerable species went extinct earlier — we found no such relationship (Fig. 7d). However, the comparison of susceptibility under increasing intensities of egg harvesting revealed the highest demographic risk from this type of activity for the extinct *Genyornis* compared to the extant *Dromaius* and *Alectura* birds, supporting the notion that egg harvesting (and not hunting of adults) might have been at least partially responsible for the demise of *Genyornis* [35]. We can also reject the hypothesis that the largest, and therefore the most physiologically buffered and mobile species, were the most resilient given the lack of relationship with the inferred extinction chronology (Fig. 2, 7d). Neither did the species with the highest sensitivities to reductions arising from the various perturbation scenarios succumb earlier (Fig. 5, 6d).

This lack of relationship to the chronology, combined with the result that many of the extant species had, in fact, some of the highest extinction susceptibilities, suggest that no obvious demographic trends can explain the great Sahul mass extinction event of the Late Pleistocene. This opens the possibility that the chronology instead reflects random extinctions, or that species succumbed to circumstantial combinations of stressors depending on the local perturbations experienced by particular populations [3, 39]. However, this conclusion does not accord well with the notion that Late Pleistocene megafauna extinctions were non-random and occurred at a much higher pace than background extinction rates [9, 31].

Another possibility is that in the case of human hunting, preferences for selecting or avoiding particular species, such as targeting larger species for more efficient returns [40], could have overridden or interacted with intrinsic demographic susceptibility. In addition, specific behavioural adaptations could potentially have rendered demographically high-risk species in fact *less* vulnerable to human hunting, such as the behavior of *Vombatus* to dig and defend burrows that were possibly too small to access by humans compared to larger burrowing or non-burrowing vombatiformes. In the case of the macropodiformes, interspecific variation in the type of locomotion — a trait not captured by our demographic models — could have contributed more to their relative susceptibility to human hunters. For example, the ability to hop at high velocities as in *Osphranter rufus* could have given it an escape advantage over the relatively slower sthenurine macropodiformes that likely employed more bipedal striding than hopping [41].

Although marsupials are widely included in studies estimating the types of mammalian demographic relationships like those we used here [42–44], more explicit consideration of their reproductive differences compared to placental mammals beyond the corrections we were able to make might further improve the resolution of future models. In particular, marsupials are born at the extreme altricial stage and complete most of their development *ex utero* through lactation [45]. This might also change the way the cost of reproduction is borne, because the unusually long period of marsupial lactation can reduce the cost of raising offspring per unit time [46, 47].

Our models, while age-structured, stochastic, and incorporating compensatory density feedback, are still simplified expressions of a species’ particular ecological and environmental contexts. As stated, our models are aspatial, yet we know that spatial processes are correlated with local extinctions across the landscape [3]. It is therefore plausible that more localized measures of extinction risk, timing, and particular climate and habitat contexts (see Appendix S8 for an examination of demographic susceptibility relative to hindcasted climate trends) could reveal subtler demographic processes at work [48]. However, Sahul’s fossil record is still generally too sparse at a regional level to test this properly [49, 50], nor do we have data indicating how spatial variation might have altered local expressions of demographic rates in long-extinct species.

Our models also ignore biotic dependencies such as predator-prey, plant-herbivore, and competition relationships that could have modified relative susceptibility in different ways depending on the community in question [51, 52]. Trophic community networks constructed for south-eastern Sahul show that bottom-up processes most strongly affect lower trophic levels, with their influence diminishing at higher trophic levels, although extinct carnivores were more vulnerable to coextinction than extant carnivores [53].

The particulars of the *Genyornis* extinction are also still debatable given the possibility that the egg shells used to date the species [54] are potentially confounded with an extinct *Progura* megapode [55, 56]. However, removing ‘*Genyornis*’ from the extinction chronology makes no difference to our overall conclusions, but it is problematic for comparing relative extinction risk between the extinct *Genyornis* and the extant *Dromaius* and *Alectura*. In fact, by including an extant megapode (*Alectura*) in our model simulations, we determined that this much smaller (2.2 kg) and faster-reproducing species had a much lower extinction susceptibility than both *Genyornis* and *Dromaius*.

That we found no clear patterns among the extinct megafauna of Sahul to explain their relative susceptibility agrees with previous work in mammals that shows that risk can be high across all body masses depending on a species’ particular ecology [57]. By definition, the megafauna were generally large (> 45 kg) species, yet their body mass does not seem to impart any particular demographic disadvantage that can explain the extinction chronology in Sahul. Our approach also provides a template for assessing relative demographic susceptibility to extinction for species in other continents that could reveal previously underappreciated dynamics and drivers, even though more spatially and community-dependent models are still needed to provide a more complete picture of the dynamics of Late Pleistocene megafauna extinctions.

## Materials and Methods

### Choice of species

Our first step was to choose enough extinct and extant species from the Sahul fossil record [49, 50] to represent a diversity of clades that were particularly affected during the main extinction event (estimated between 60 and 40 ka, with 1 ka = 1000 years ago) [58]. We also aimed to include at least one extant species within each functional/taxonomic group to compare extant with extinct species’ susceptibility. We settled on a total of twenty-one species (13 extinct; 8 extant) from five different functional/taxonomic groups: (***i***) **five vombatiform herbivores**, (***ii***) **seven macropodiform herbivores**, (***iii***) **three large birds**, (***iv***) **four carnivores**, and (***v***) **two monotreme invertivores**. For a full list and justification of species chosen as well as the distribution of mean body masses, refer to Supplementary Information Appendix S1.

### Estimating demographic rates

To build age-structured population models for extinct taxa, we relied on different allometric, phylogenetic, and measured relationships to predict the plausible range of component demographic rates. For most extant marsupials, we relied mainly on the marsupial life-history database published in Fisher *et al*. [59], but updated some values for some species with more recent references (see below). A detailed description of how we estimated the necessary demographic rates and other ecological data to build the stochastic models is provided in Supplementary Information Appendix S2, and a full table of all demographic values is provided in Appendix S2, Table S1).

### Age-structured (Leslie) population models

From the estimated demographic rates for each species, we constructed a pre-breeding, *ω*+1 (*i*) × *ω*+1 (*j*) element (representing ages from 0 to *ω* years old), Leslie transition matrix (**M**) for females only (males are demographically irrelevant assuming equal sex ratios). Fertilities (*m_x_*) occupied the first row of the matrix, survival probabilities (*S_x_*) occupied the sub-diagonal, and we set the final diagonal transition probability (**M**_*ij*_) to *S_ω_* for all species except *Vombatus, Thylacinus* and *Sarcophilus* for which we instead set to 0 to limit unrealistically high proportions of old individuals in the population, and the evidence for catastrophic mortality at *ω* for the latter two species (dasyurids) [60–62]. Multiplying **M** by a population vector **n** estimates total population size at each forecasted time step [63]. Here, we used **n**_0_ = *AD***Mw**, where **w** = the right eigenvector of **M** (stable stage distribution), and *A* = the surface area of the study zone applied in the stochastic extinction scenarios — we arbitrarily chose *A* = 250,000 km^2^ (500 km × 500 km; approximately 10% larger than the state of Victoria) so that the species with the lowest **n**_0_ would have a population of at least several thousand individuals at the start of the simulations (see Supplementary Information Appendix S3). We also included a compensatory density-feedback function in all simulations to avoid exponentially increasing populations (see Supplementary Information Appendix S4).

### Stochastic extinction scenarios

With the base **M** including density feedback tailored for each species, we perturbed various elements of their life histories to test hypotheses regarding plausible extinction drivers and pathways (see Fig. 1). We first tested the relationship between extinction date and speed of life history as a baseline without any perturbation (Scenario *i*), and then we generated six additional scenarios with perturbations (Scenarios *ii–vii*). The second scenario increased juvenile (*x* = 0 to *α*-1) mortality (plus a sub-scenario [*iib*] where we progressively removed individual juveniles from the population as we did for all individuals in Scenario *v* — see below). This scenario aims to emulate either food shortages of sufficient magnitude to make growing juveniles with higher relative energy and water demand [32] succumb to environmental change more than adults, or from targeted hunting of juveniles by humans [25]. The third scenario progressively reduces fertility to emulate food shortages lowering energetically demanding reproduction/lactation [33, 34]. We also considered a sub-scenario (Scenario *iiib*, see details below) for the category of large birds where we progressively increased the number of eggs harvested by humans [35]. In Scenario *iv*, we progressively reduced survival across all age classes to examine the influence of an age-independent environmental stressor. Scenario *v* progressively removed individuals from the **n** population vector emulating offtake where animals are directly removed from the population to simulate human hunting (with age-relative offtake following the stable stage distribution of the target species). In Scenario *vi*, we emulated how environmental variability would compromise populations via an increased relative (i.e., per generation) frequency of catastrophic die-offs by progressively increasing the number of catastrophic ~ 50% mortality events occurring per generation. Finally, Scenario *vii* progressively increased the magnitude of the catastrophic mortality events to examine species’ responses to rising severity of catastrophes[36].

For Scenario *iiib*, we estimated the egg-production component for *Genyornis* by calculating the proportion of total fecundity contributed by individual egg production in *Dromaius* (nest success of 0.406 × hatching probability of 0.419 = 0.17), and then multiplying this proportion by the total fertility estimated for *Genyornis* from equation 11 — this produced an estimated per-individual annual egg production of 7.74 eggs for *Genyornis* (or, 7.74/2 = 3.87 eggs resulting in daughters). For Scenario *vii*, we randomly allocated a multiplier of the expected frequency per generation (uniform sampling) derived from the species-specific range of multipliers identified in Scenario *vi* (i.e., from 1 to the value where the species has an extinction probability = 1). In this way, we both standardized the relative risk among species and avoided cases where catastrophe frequency was insufficient to elicit any iterations without at least one extinction.

We ran 10,000 stochastic iterations of each model starting with allometrically predicted stable population size (see Supplementary Information Appendix S3) divided into age classes according to the stable stage distribution. We projected all runs to 40[*G*] for each species (removing the first [*G*] values as burn-in). In each scenario, we progressively increased the relevant perturbation and calculated the proportion of 10,000 stochastic model runs where the final population size fell below a quasi-extinction (E_q_) of 50 female individuals (100 total individuals total assuming 1:1 sex ratios). This threshold is based on the updated minimum size below which a population cannot avoid inbreeding depression [64]. After calculating the per-increment probability of E_q_ in each of the seven scenarios, we calculated the total area under the quasi-extinction curve (integral) for each species as a scenario-specific representation of extinction risk across the entire range of the specific perturbation — this provides a single, relative value per species for comparison. Finally, we ranked the integrals among species in each scenario (lower ranks = higher resilience), and took the median rank as an index of resilience to extinction incorporating all scenario sensitivities into one value for each species.

### Extinction dates

We compared the relative susceptibilities among all extinction scenarios, as well as the combined extinction-susceptibility rank of each species, to estimates of continental extinction times for the genera we examined. We took all estimates of continental extinction dates from the Signor-Lipps corrected values provided in Saltré *et al*. [58]; however, more recent continent-wide disappearance dates for *Thylacinus* and *Sarcophilus* are provided in White *et al*. [38].

We hypothesize that one, or several, of these types of perturbation (Scenarios *ii–vii*) would lead to a better match between the continental-scale chronology of extinctions as inferred from the fossil record compared to the simpler expectation of larger species with slower life-histories being more likely to go extinct than smaller species with faster life histories when faced with novel mortality sources (Scenario *i*) [37].

## Data and code availability

All data and are R code needed to reproduce the analyses are available for download at github.com/cjabradshaw/MegafaunaSusceptibility.

## Acknowledgments

This study was supported by the Australian Research Council through a Centre of Excellence grant (CE170100015) to C.J.A.B., C.N.J., and V.B., and a Discovery Project grant (DP170103227) to V.B. We acknowledge the Indigenous Traditional Owners of the land on which Flinders University is built — the Kaurna people of the Adelaide Plains.

## Supplementary Information

Contents:

**Appendix S1**. Choice of species and body mass distribution (includes Fig. S1)

**Appendix S2**. Estimating demographic rates and population data as input parameters for the stochastic models (includes Table S1)

**Appendix S3**. Allometric predictions of equilibrium population density, total population size, and biomass (includes Fig. S2)

**Appendix S4**. Compensatory density feedback

**Appendix S5**. Deriving marsupial correction factors for fecundity (*F*) and age at first breeding (*α*) (includes Fig. S3–S6)

**Appendix S6**. Quasi-extinction curves for each species in each of the 6 perturbation scenarios (Scenarios *ii—vii*) (includes Fig. S7)

**Appendix S7**. Comparing increasing mortality in juveniles and increasing offtake of juvenile individuals (Includes Fig. S8)

**Appendix S8**. Comparing demographic susceptibility to climate variation (includes Fig. S9)

**Supplementary Information References**

## Appendix S1. Choice of species and body mass distribution

Given data availability, we settled on a total of twenty-one species (13 extinct; 8 extant) from five different functional/taxonomic groups: (***i***) **five vombatiform herbivores**: *Diprotodon optatum* (2786 kg; extinct) [1], *Palorchestes azael* (1000 kg; extinct) [2], *Zygomaturus trilobus* (500 kg; extinct) [3], *Phascolonus gigas* (200 kg; extinct) [3], and *Vombatus ursinus* (common wombat; 25 kg; extant) [4, 5]; (***ii***) **seven macropodiform herbivores**: *Procoptodon goliah* (250 kg; extinct) [6], *Sthenurus stirlingi* (150 kg; extinct) [6], *Protemnodon anak* (130 kg; extinct) [3], *Simosthenurus occidentalis* (120 kg; extinct) [3], *Metasthenurus newtonae* (55 kg; extinct) [3], *Osphranter rufus* (red kangaroo; 25 kg; extant) [7], and *Notamacropus rufogriseus* (red-necked wallaby; 14 kg; extant) [8]; (***iii***) **three** omnivorous (but primarily plant-eating) **large birds**: *Genyornis newtoni* (200 kg; extinct) [3], *Dromaius novaehollandiae* (emu; 55 kg; extant) [9], and *Alectura lathami* (brush turkey; 2.2 kg; extant) [10]; (***iv***) **four carnivores**: *Thylacoleo carnifex* (marsupial ‘lion’; 110 kg; extinct) [3], *Thylacinus cynocephalus* (marsupial ‘tiger’; 20 kg; extinct) [11], *Sarcophilus harrisii* (devil; 6.1 kg; extinct in mainland Australia, but extant in Tasmania — see below) [12], and *Dasyurus maculatus* (spotted-tail quoll; 2.0 kg; extant) [13]; and (***v***) **two monotreme invertivores**: *Megalibgwilia ramsayi* (11 kg; extinct) [3], and *Tachyglossus aculeatus* (short-beaked echidna; 4.0 kg; extant) [14].

For each species, we identified the body mass of mature females. However, the sex of an extinct individual from its fossilized remains in many species is difficult to determine, especially when sample sizes are small [15]. As such, we might have inadvertently assigned a female mass based on an estimated male mass, given evidence that there is a male bias in many fossil collections [16, 17]. For this reason, we attempted to cover a broad range of body masses among species to maximize the *relative* difference between them for comparison.

The two genera *Thylacinus* and *Sarcophilus* require special consideration in both the design of the analysis and the interpretation of the results. While *Sarcophilus* is extant, it is restricted to the island state of Tasmania that has been separated from mainland Australia since approximately 8–10 ka. However, the species went extinct on the mainland 3179 (± 24) years ago, whether considering the species complex *Sarcophilus harrisii* and *S. laniarius* together or separately [18]. Like *Sarcophilus, Thylacinus* went extinct on the mainland just over 3000 years ago [18] and persisted in Tasmania. However, unlike *Sarcophilus, Thylacinus* also went extinct in Tasmania in the 1930s. In our analyses we treat *Sarcophilus* as ‘extant’, with the proviso that it should be considered extinct on the mainland. Although we could also have treated *Thylacinus as* ‘extant’ in the sense that it persisted into historical times in Tasmania, we treat this species as extinct in our analyses.

The distribution of the body masses (*M*) across the nineteen species (range: 1.68–2786 kg) was approximately log-Normal (Shapiro-Wilk Normality test on log_10_*M*: *W* = 0.9804; *p* = 0.9305; Fig. S1). Median dates of continental (i.e., total species) extinction ranged from 51,470 (± 3167 standard deviation) for *Metasthenurus*, to 3179 (± 24) for *Sarcophilus* (mainland only; currently extant in Tasmania) [18, 19].

**Figure S1.**
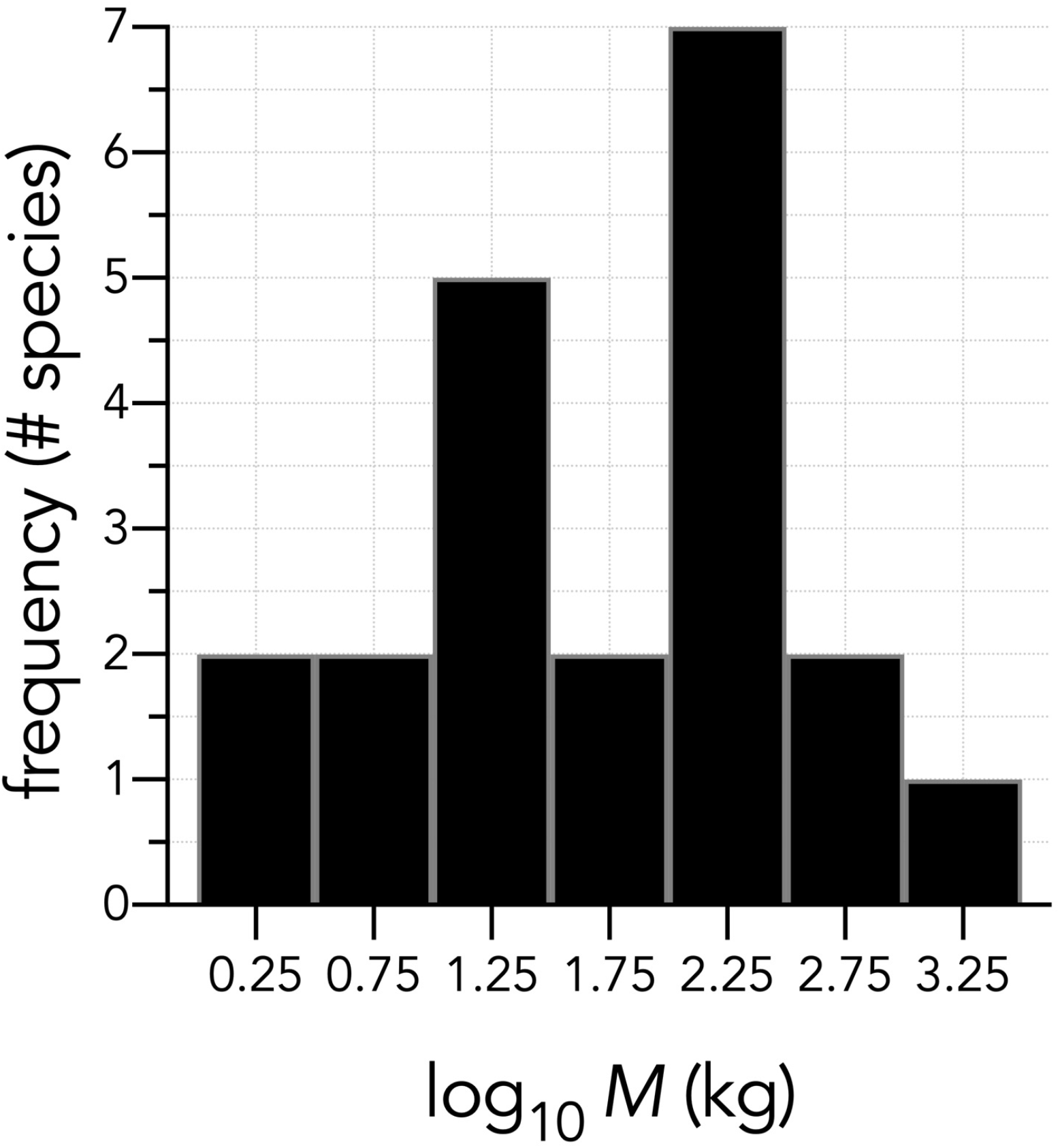
Histogram of log_10_ adult body masses (in kg) for the twenty-one species examined. The distribution is approximately log-Normal (Shapiro-Wilk Normality test on log_10_*M*: *W* = 0.9804; *p* = 0.9305).

## Appendix S2. Estimating demographic rates and population data as input parameters for the stochastic models

For each species, we first calculated the maximum rate of instantaneous population growth (*r_m_*) using the following equation:

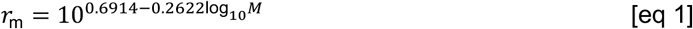

for mammals, where *M* = mass (g) [20]. For birds, we used an optimization of the objective function based on age at first breeding (*α*, estimated as shown below), adult survival (*s*_ad_, estimated as shown below), and the allometric constant for birds [21] *a_rT_* = 1.107:

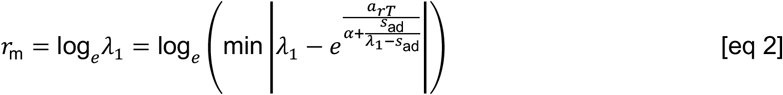

We then calculated theoretical equilibrium population densities (*D*, km^-2^) based on the following:

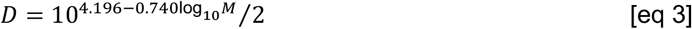

for mammalian herbivores (*M* = body mass in g), where dividing by 2 predicts for females only (i.e., assumed 1:1 sex ratio) [22]. For large, flightless birds (i.e., *Genyornis* and *Dromaius*) [23], we applied:

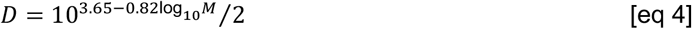

and

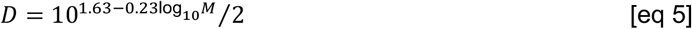

for omnivorous birds (i.e., *Alectura*) [24] where *M* is in g. For mammalian carnivores, we applied:

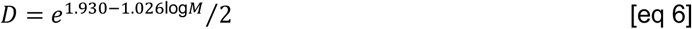

(*M* in kg), which we derived from Stephens *et al*. [25]. There were no specific invertivore or taxonomically specific equations to estimate *D*; however, we determined that the equation for the fitted 97.5 percentile in mammalian carnivores:

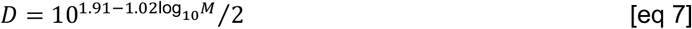

Stephens *et al*. [25] provided is a reasonable *D* for female *Tachyglossus* = 9.9 km^-2^. This is comparable to echidna densities measured for Kangaroo Island (4.4 females km^-2^) [26] and Tasmania (8.4 females km^-2^) [14]. We therefore also used equation 7 to predict *D* for *Megalibgwilia*. For a detailed description of the distribution and trends of equilibrium densities, population sizes, and biomasses across the modelled species, see Appendix S3.

We estimated the maximum age (*ω*) of each species according to:

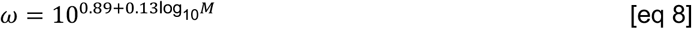

for non-volant birds and mammals (*M* in g) [27], or

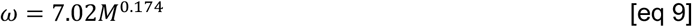

for *Alectura* (*M* in g) [28], the latter of which produces a *ω* that closely matches the maximum longevity of 25 years estimated for the similar-sized megapode *Leipoa ocellata* (malleefowl) [29].

For other species, we made species- or group-specific adjustments to the estimates of *ω*: for *Vombatus* [7], we set *ω* = 26; the disparity between this and the *ω* derived from the allometric prediction (equation 7 gives *ω* = 29) means we adjusted *ω*′ = 26/29*ω* for all vombatiform herbivores. Similarly for the macropodiformes, we scaled the predicted *ω* according to the degree of overprediction of the parameter from the equation for *Osphranter rufus* (for the latter, the equation predicted *ω* = 30, but in reality it is closer to 13) [30], meaning we adjusted *ω*′ = 13/30*ω* for all macropodiformes except *Notamacropus*; for *Notamacropus rufogriseus*, we took the mean maximum age[31] of 16. For *Dromaius* [32], we set *ω* = 17; for *Thylacoleo*, we set *ω* = 17 to match female lion *Panthera leo* longevity [33]; for *Thylacinus* [34, 35], we set *ω* = 10, for *Sarcophilus* [36] *ω* = 5, and *Dasyurus ω* = 4 given the catastrophic mortality at maximum lifespan characteristic of dasyurids [37–39]. In the case of *Dasyurus* (both *D. maculatus* and *D. hallucatus*), most females die in their third year, although some can persist into the fourth year [40–43] and maximum longevity in captivity can be up to 5 years [44]. For *Tachyglossus, ω* is extremely high compared to similar-sized marsupials or placentals: up to 50 years in captivity, and possibly 45 years in the wild [14]; we set the latter value of *ω* = 45 to be conservative. For *Megalibgwilia*, we assumed that the underestimation for *Tachyglossus* according to equation 8 would also apply, so here we set *Megalibgwilia ω* = (45/23)26 = 51 years.We estimated fecundity (*F*; mean number of female neonates produced per year and per breeding female) for mammals [45] as:

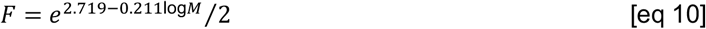

dividing by 2 for daughters only (*M* in g). Although we used well-establish allometric relationships to derive our input parameters, most of these relationships are based on placental species. It is accepted that average life history traits differ between similar-sized marsupials and placentals [46], and we therefore estimated a correction factor for *F* and age at first breeding (*α*) (see equation 12) for both the vombatiform and macropodiform herbivore groups separately (see Appendix S5, Fig. S3–S5 for approach) using demographic data describing marsupial life histories [46].

For *Vombatus* [5], we set *F* = 0.25 given a litter size = 1, an inter-birth interval of 2 years, and an assumed sex ratio of 1:1, from which we derived *F* for the extinct vombatiform herbivores (Appendix S5, Fig. S4). For *Notamacropus rufogriseus*, we set the average annual number of offspring = 1 multiplied by a 2.8% twinning rate [47], 1.3 based on an interbirth interval of 286 days, and an assumed 1:1 sex ratio. For *Genyornis*, we applied the following equation:

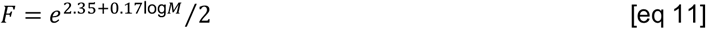

(*M* in g) [45]. For *Dromaius* [9], we used the average of 6.7 eggs clutch^-1^ and 3.4 clutches year^-1^, an assumed sex ratio of 1:1, and nest success (0.406) and hatching probabilities (0.419) for ostriches [48]. For *Alectura*, we used the annual mean of 16.6 eggs pair^-1^ for *Leipoa ocellata* [29], a hatching success of 0.866 for *Alectura lathami* [49], and an assumed 1:1 sex ratio. For *Thylacoleo*, we applied the average litter size of 1 for large vombatiforms, and a 1:1 sex ratio; for *Thylacinus* and *Sarcophilus*, we applied the values of 3.42 progeny litter^-1^ and the proportion [35, 50] of adults reproducing year^-1^ (0.91), and a 1:1 sex ratio. The allometric prediction (equation 10) nearly matched the product of mean litter size (4.9) [40] and proportion females breeding (0.643) [41] for *Dasyurus*, so we used the former. For *Tachyglossus* [51], we used the production of 1 egg breeding event (divided by 2 for daughters) multiplied by the probability of breeding = 0.55. For *Megalibgwilia*, we assumed that the overestimation for *Tachyglossus* according to equation 10 would also apply, so here we set *Megalibgwilia F* = (0.275/0.659)0.532 = 0.222.

To estimate the age at first breeding (*α*), we used the following relationship for mammals [28]:

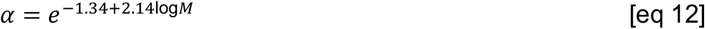

and

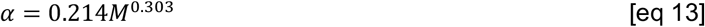

for birds [28] (*M* in g). For the macropodiforms, equation 12 appeared to overestimate *α* by ~ 20% (see Appendix S5, Fig. S4), so we adjusted the extinct macropodiforms accordingly. For the vombatiform herbivores, equation 12 performed more according to expectation. For example, the 1800-2000 kg female white rhinoceros (*Ceratotherium simum*) has *α* = 6–7 years [52], which is similar to the 7 years predicted by equation 12 for the 2786 kg *Diprotodon optatum* (but < *α* = 10–12 years for the > 6000 kg female African savanna elephant *Loxodonta Africana*) [53]. We also made species-specific adjustments to *α* for the extant species (or recently extinct in the case of *Thylacinus*). In the case of *Vombatus* [54], we set *α* = 2, *α* = 2 for *Osphranter rufus* [30], *α* = 1 for *Notamacropus rufogriseus* [47], *α* = 3 for *Dromaius* [55], *α* = 2 for *Alectura* based on data from *Leipoa ocellata* [29, 56], *α* = 1 for *Thylacinus* [35, 50], *Sarcophilus* [36], and *Dasyurus* [41]. For *Thylacinus* and *Sarcophilus*, although *α* = 1, only a small proportion of females breed at this age (see below), so for most females *α* is in fact 2. But for *Dasyurus*, some females can become sexually mature and breed at < 12 months old [43], so we incorporated a modest capacity to reproduce in the year prior to age 1 (40% of total fertility). For *Tachyglosssus*, we set *α* = 3 based on evidence that echidnas take 3-5 years to reach adult mass [14], and only adults are observed to breed [26]; as such, we set *m* = 0.5*F* in year 3, 0.75*F* in year 4, and *m* = *F* thereafter. As we did for *ω* and *F*, we estimated the bias between the allometrically predicted and measured *α* (equation 12) for *Tachyglossus*, and applied this to *Megalibgwilia;* however, rounding to the nearest year also means *α* = 3 for *Meglibgwilia*. The log_10_ of the resulting *α* among species predicted their respective log_10_ *r*_m_ well (*R^2^* = 0.73) (Appendix S5, Fig. S6), with the fitted parameters similar to theoretical expectation [57], and thus, supporting our estimates of *r*_m_ as realistic.

To estimate age-specific fertilities (*m_x_*) from *F* and *α*, we fit a logistic power function of the general form:

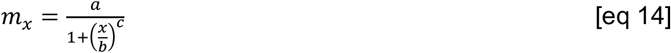

where *x* = age in years, and *a*, *b*, *c* are constants estimated for each species, to a vector composed of (*α*-1) values at 0*F*, [*α*/2| values at 0.75*F*, and for the remaining ages up to *ω*, the full value of *F*. This produced a continuous increase in *m_x_* up to maximum rather than a less-realistic stepped series. For *Sarcophilus*, we instead used the parameters from an existing devil model [36] to populate the *m_x_* vector.

To estimate realistic survival schedules, we first used the allometric prediction of adult survival (*s*_ad_) as:

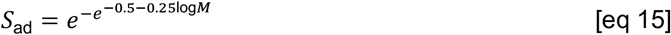

for mammals, and:

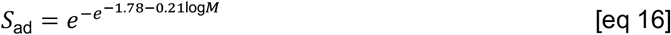

for birds, where *M* = body mass (g) [58]. For *Tachyglossus*, we used the mean *S*_ad_ = 0.96 based on upper and lower estimates of mortality for tagged individuals over 15 years in Tasmania [51], and applying the allometric-bias correction for *Megalibgwilia* as for *ω*, *F*, and *α* as described above. We then applied the Siler hazard model [59] to estimate the age- (*x*-) specific proportion of surviving individuals (*l_x_*); this combines survival schedules for immature, mature, and senescent individuals within the population:

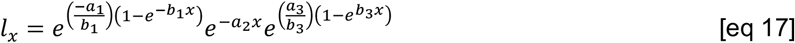

where *a*_1_ = initial immature mortality, *b*_1_ = rate of mortality decline in immatures, *a*_2_ = the ageindependent mortality due to environmental variation, *a*_3_ = initial adult mortality, and *b*_3_ = the rate of mortality increase (senescence). From *l_x_*, age-specific survival can be estimated as:

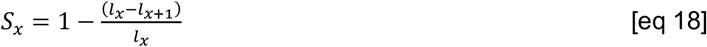

We estimated the component parameters starting with 1 - *S*_ad_ for *a*_1_ and *a*_2_, adjusting the other parameters in turn to produce a dominant eigen value (*Λ*i) from the transition matrix containing *S_x_* (see *Methods*) such that log_*e*_ λ_1_ ≈ *r*_m_. However, in many cases, marsupial and monotreme life histories were incapable of reproducing predicted *r*_m_ (see Table S1 below), although we attempted to maximize log_*e*_ Λ_1_ wherever possible. This appears to be biologically justified given the slower life histories of vombatiforms and monotremes in particular compared to macropodiforms. We also generally favoured a more pronounced senescence component of *l_x_* in the longer-lived species given evidence for survival senescence in long-lived mammals [60]. For *Sarcophilus*, we instead used the parameters from an existing devil model [36] to populate the *S_x_* vector.

**Table S1.**
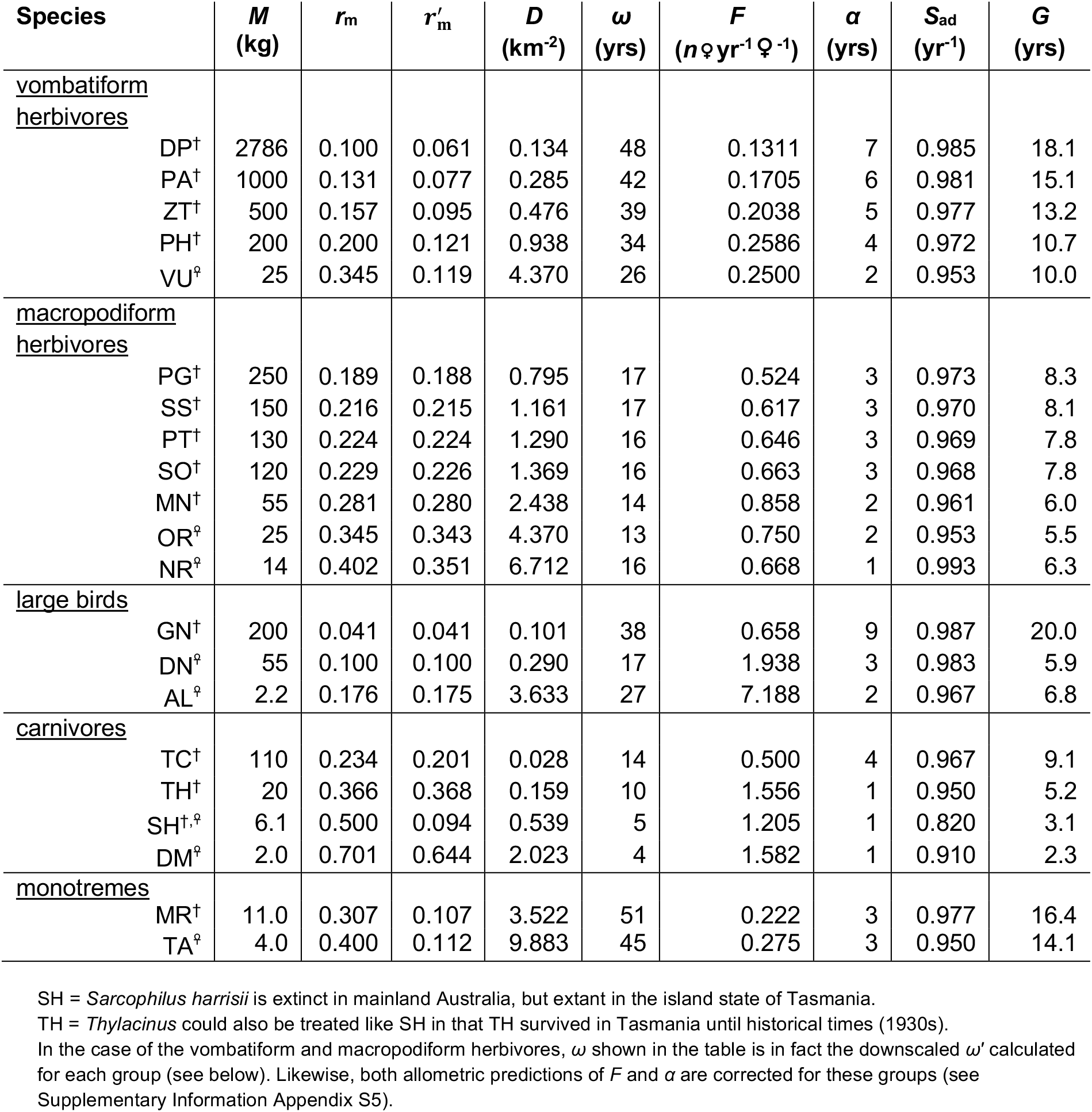
Predicted demographic values for each species (equations provided in *Methods*). *M* = mass, *r*_m_ = maximum rate of instaneous exponential population growth predicted allometrically, 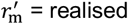 *r*_m_ predicted from the constructed matrix (see text), *ω* = longevity, *F* = fertility (daughters per breeding female per year), *α* = age at first reproduction (primiparity), *S*_ad_ = yearly adult survival, *G* = generation length. Species notation: DP = *Diprotodon optatum*, PA = *Palorchestes azael*, ZT = *Zygomaturus trilobus*, PH = *Phascolonus gigas*, VU *Vombatus ursinus* (vombatiform herbivores); PG = *Procoptodon goliah*, SS = *Sthenurus stirlingi*, PT = *Protemnodon anak*, SO = *Simosthenurus occidentalis*, MN = *Metasthenurus newtonae*, OR = *Osphranter rufus*, NR = *Notamacropus rufogriseus*, common name: red-necked wallaby (macropodiform herbivores); GN = *Genyornis newtoni*, DN = *Dromaius novaehollandiae*, AL = *Alectura lathami* (large birds); TC = *Thylacoleo carnifex*, TH = *Thylacinus cynocephalus*, SH = *Sarcophilus harrisii*, DM = *Dasyurus maculatus* (carnivores); TA = *Tachyglossus aculeatus*, MR = *Megalibgwilia ramsayi* (monotreme invertivores). ^†^extinct; ^♀^extant.

## Appendix S3. Allometric predictions of equilibrium population density, total population size, and biomass

The allometric predictions of stable population size (*N*_stable_) for each species in the 500 km × 500 km study area showed the largest populations for some of the smallest, extant species (e.g., *N*_stable_ > 600,000 for *Vombatus, Osphranter, Notamacropus, Alectura, Dasyurus, Tachyglossus*) (Fig. S2a). There was a clear separation in the allometric predictions of *N*_stable_ among the species in each group (Fig. S2b). When expressed as total biomass across the study area, the four carnivores had approximately equal biomasses (~ 10^6^ kg), as did the macropodiformes and monotremes (Fig. S2c). For the large birds and herbivore vombatiformes, biomass increased with body mass (Fig. S2c).

**Figure S2.**
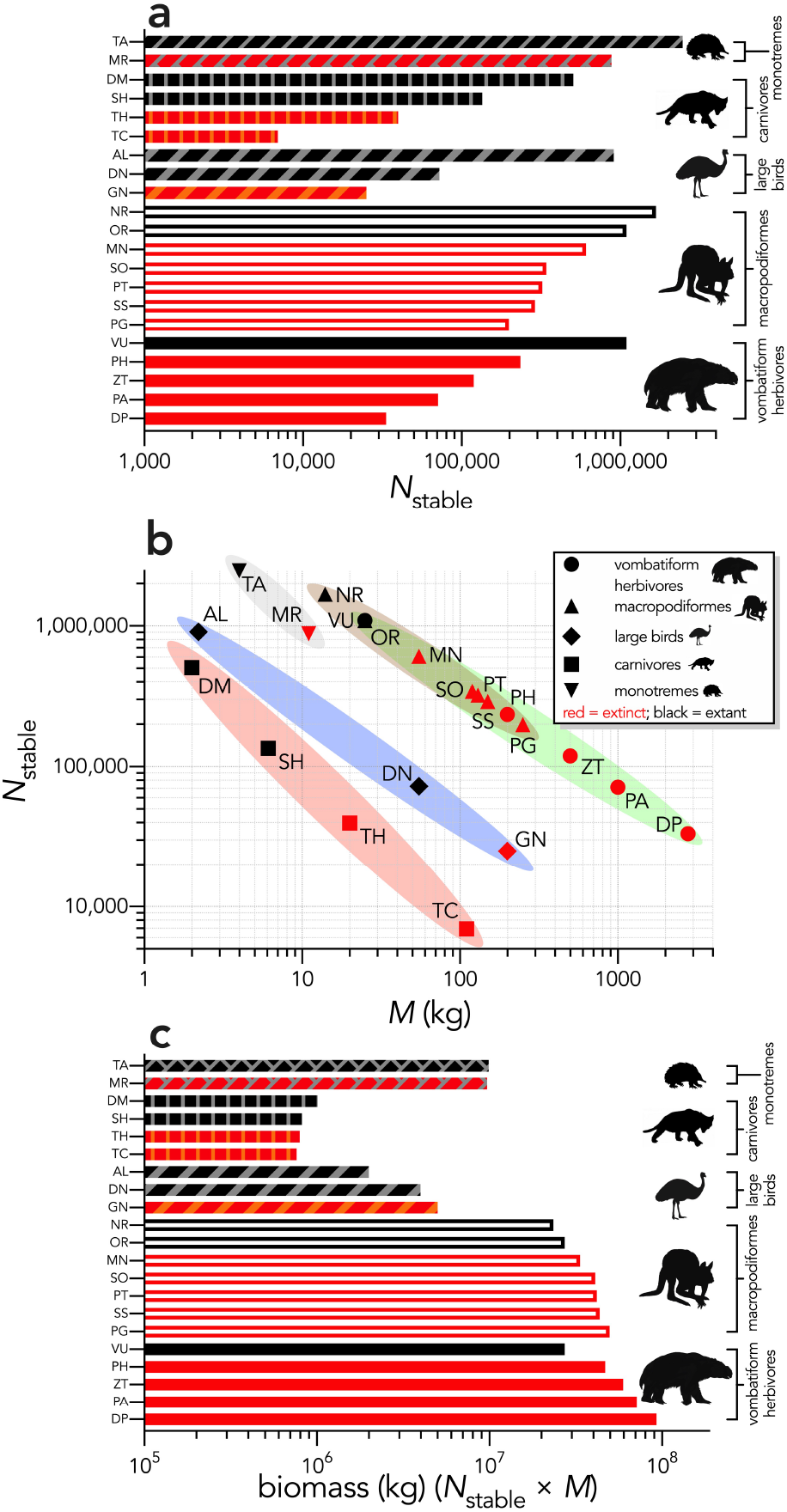
(a) Stable population sizes (*N*_stable_) for each modelled species predicted from allometric estimates of population density for a 500 km × 500 km (250,000 km^2^) landscape; (b) *N*_stable_ plotted against body mass (*M*, in kg), showing the allometric scaling separating the vombatiform herbivores (green)/macropodiformes (brown), flightless birds (blue), carnivores (red), and monotremes (grey); (c) predicted landscape biomass (*N*_stable_ × *M*) for each species. Species notation: DP = *Diprotodon optatum*, PA = *Palorchestes azael*, ZT = *Zygomaturus trilobus*, PH = *Phascolonus gigas*, VU *Vombatus ursinus* (**vombatiform herbivores**); PG = *Procoptodon goliah*, SS = *Sthenurus stirlingi*, PT = *Protemnodon anak*, SO = *Simosthenurus occidentalis*, MN = *Metasthenurus newtonae*, OR = *Osphranter rufus*, NR = *Notamacropus rufogriseus*, common name: red-necked wallaby (**macropodiformes**); GN = *Genyornis newtoni*, DN = *Dromaius novaehollandiae*, AL = *Alectura lathami* (**large birds**); TC = *Thylacoleo carnifex*, TH = *Thylacinus cynocephalus*, SH = *Sarcophilus harrisii*, DM = *Dasyurus maculatus* (**carnivores**); TA = *Tachyglossus aculeatus*, MR = *Megalibgwilia ramsayi* (**monotreme invertivores**). Here, we have depicted SH as ‘extant’, even though it went extinct on the mainland > 3000 years ago.

## Appendix S4. Compensatory density feedback

To avoid an exponentially increasing population without limit generated by a transition matrix optimized to produce values as close to *r*_m_ as possible, we applied a theoretical compensatory density-feedback function. This procedure ensures that the long-term population dynamics were approximately stable by creating a second logistic function of the same form as *m_x_* to calculate a modifier (*S*_mod_) of the *S_x_* vector according to total population size (Σ**n**):

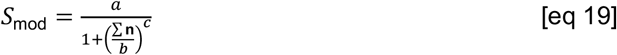

We adjusted the *a*, *b*, and *c* constants for each species in turn so that a stochastic projection of the population remained stable on average for 40 generations (40[*G*]), where:

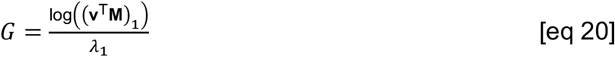

and (**v**^⊤^**M**)_1_ = the dominant eigen value of the reproductive matrix **R** derived from **M**, and **v** = the left eigenvector[61] of **M**. Although arbitrary, we chose a 40[G] projection time as a convention of population viability analysis to standardize across different life histories [62, 63].

The projections were stochastic in that we *β*-resampled the *S_x_* vector assuming a 5% standard deviation of each *S_x_* and Gaussian-resampled the *m_x_* vector at each yearly time step to 40[*G*]. We also added a catastrophic die-off function to account for the probability of catastrophic mortality events (*C*) scaling to generation length among vertebrates [64]:

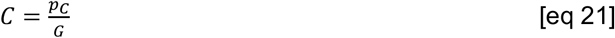

where *p_c_* = probability of catastrophe [64] (set at 0.14). Once invoked at probability *C*, we applied a *β*-resampled proportion centered on 0.5 to the *β*-resampled *S_x_* vector to induce a ~ 50% mortality event for that year [65], as we assumed that a catastrophic event is defined as any 1 yr peak-to-trough decline in estimated numbers of 50% or greater” [64]. Finally, for each species we rejected the first ⌊*G*⌋ years of the projection as a burn-in to allow the initial (deterministic) stable stage distribution to stabilize to the stochastic expression of stability under compensatory density feedback.

## Appendix S5. Deriving marsupial correction factors for fecundity (*F*) and age at first breeding (*α*)

Given that allometric predictions of various life-history traits in mammals are based primarily on data from extant placentals, we investigated the degree of potential bias in our estimates of longevity (*ω*) fecundity, (*F*) and age at first breeding (*α*) based on a comparison of theoretical and observed data for extant marsupials. We discuss the bias correction *ω* for in the main text for the vombatiform and macropodiform herbivores (see Appendix S2), so here we report how we derived group-specific corrections for *F* and *α*.

### Fertility (F) correction

We first collected adult female mass, inter-birth interval (*I*_b_), and age at first breeding data for twenty-three extant species in the database compiled by Fisher *et al*. [46]. We included all species for which data were listed in the genera: *Macropus (Osphranter* and *Notamacropus), Dorcopsis, Lagorchestes, Petrogale, Thylogale*, and *Wallabia*. We excluded the genus *Dendrolagus* (tree kangaroos) because they represent a distinct clade and differ ecologically from most other macropodids. We also excluded the Tammar wallaby (*Macropus eugenii*) because it is a strongly seasonal-breeding species that potentially strong leverage on estimating the allometric slope.

To correct *F*, we first examined the relationship between *I*_b_ and body mass (*M*) for these species (Fig. S3):

**Figure S3.**
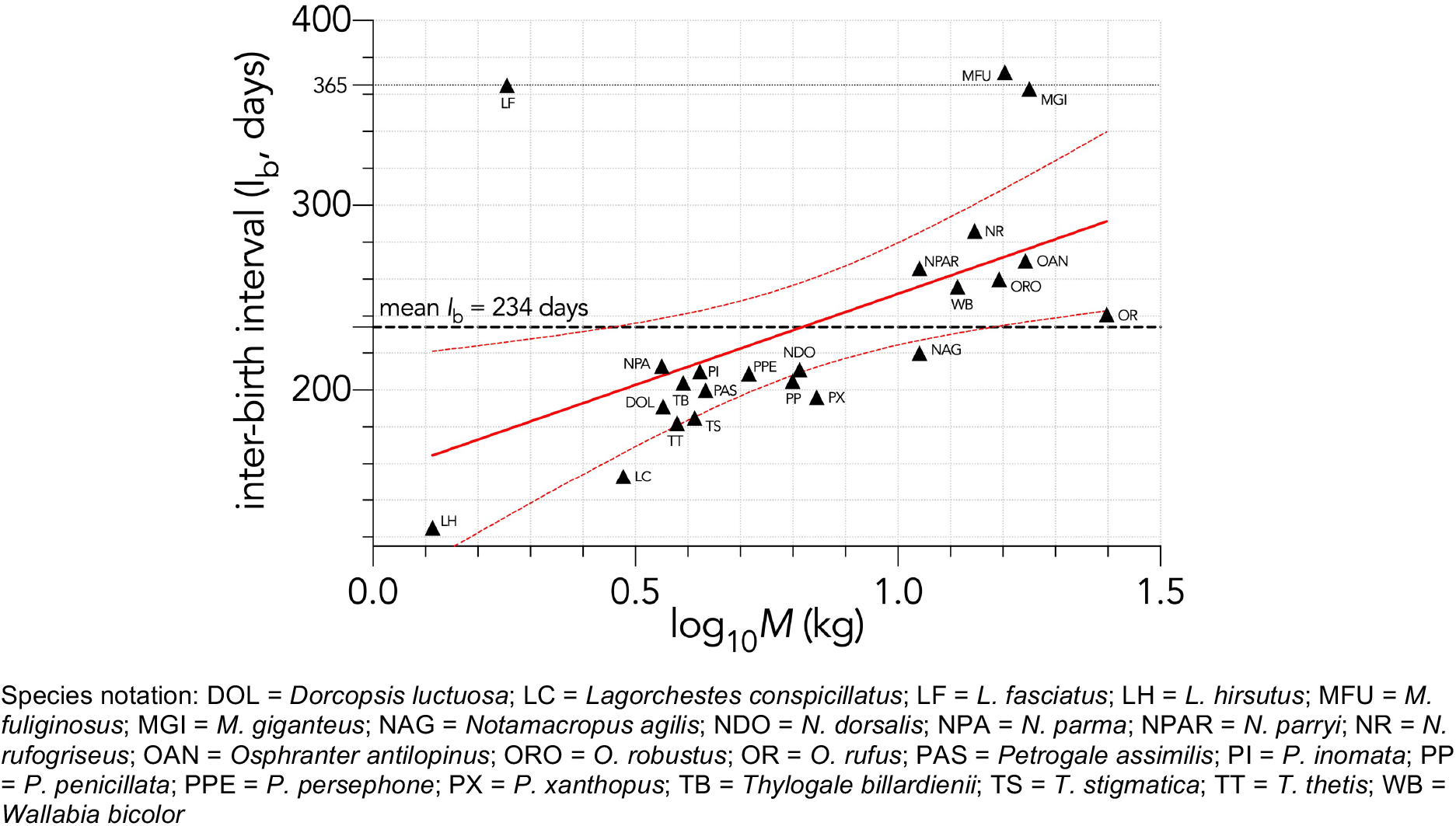
Relationship between the logarithm of adult female body mass (*M*, kg) and inter-birth interval (*I*_b_, in days) for 23 extant macropodid herbivores[46]. The estimated parameters of the linear fit (*y* ~ *α* + *βx*) are: *α* = 159.3 ± 31.5 days (± SE) and *β* = 93.6 ± 36.4, and explaining 24.2% of the variation 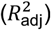, with the information-theoretic evidence ratio (ER) of the slope *versus* intercept-only model = 11.0.

We therefore concluded that there was sufficient evidence for an allometric relationship between the two variables for this group, which we used to project the degree to which *F* was overestimated by the allometric relationship (equation 10) used to estimate *F* for the extinct macropodidiform herbivores. Using the intercept and slope estimated in the relationship shown in Fig. S3, we predicted an inter-birth interval of 384 days for *Procoptodon*, 363 days for *Sthenurus*, 357 days for *Protemnodon*, 354 days for *Simosthenurus*, and 322 days for *Metasthenurus*. These changed the allometrically predicted *F* by −4.9%, +0.5%, +2.2%, +3.1%, and +13.3%, respectively (corrected *F* shown in Appendix 2, Table S1).

For the vombatiforms, there are only four extant phascolarctid (koala *Phascolarctos cinereus*) and vombatiform herbivores (common or bare-nosed wombat *Vombatus ursinus*, northern hairy-nosed wombat *Lasiorhinus krefftii*, and southern hairy-nosed wombat *L. latifrons*). There were not enough species to estimate an allometric relationship that might predict the expected *Ib* for extinct vombatiform herbivores, so instead we assumed that the extinct vombatiform herbivores we considered would scale allometrically relative to *Vombatus ursinus*, which has a measured *I*_b_ of 730 days [46]. For this, we assumed the same slope as measured for the extant macropodiforms (Fig. S3), and an intercept that aligned *Vombatus* with the relationship (Fig. S4):

**Figure S4.**
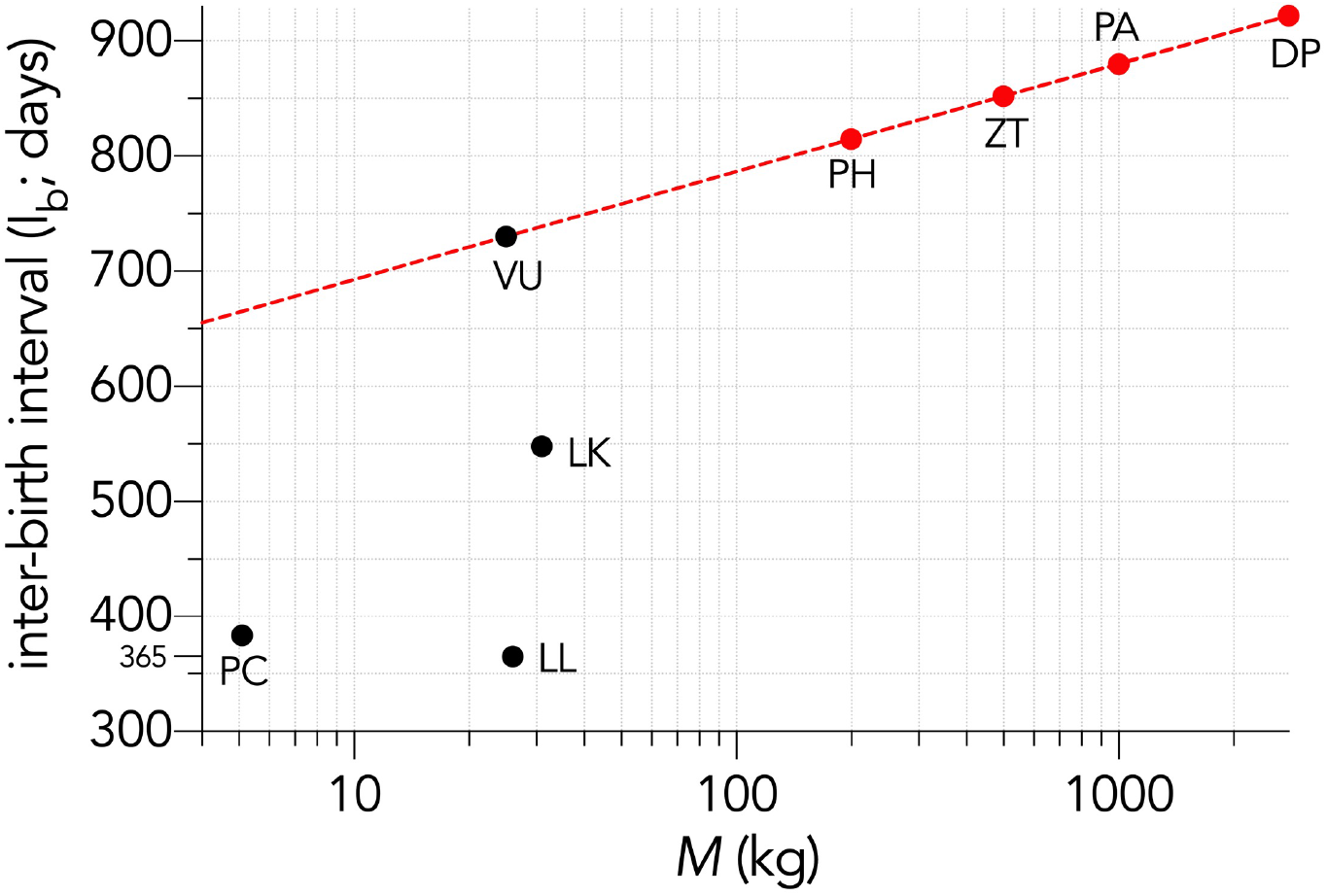
Relationship between the logarithm of adult female body mass (*M*, kg) and inter-birth interval (*I*_b_, in days) for four large, extant phoscolarctid and vombatiform herbivores: PC = *Phascolarctos cinereus;* VU = *Vombatus ursinus*; LK = *Lasiorhinus krefftii*; LL = *L. latifrons*. Shown is the assumed relationship between *I*_b_ and log_10_*M* setting the slope to that estimated for the extant macropodiforms (*β* = 93.6; Fig. S3) and an intercept that aligned with *Vombatus* (*α* = 599 days)

### Age at first breeding (α) correction

Next, we estimated the bias in the predicted age at first breeding (*α*) for the macropodiforms. A similar correction for the vombatiform herbivores was not warranted given that the allometric predictions were close to expectation for placentals of similar mass (see main text). We plotted *α* predicted from the allometric equation 12 against those observed from the marsupial database [46] for the extant macropodiform species as described above for *F* (Fig. S5):

**Figure S5.**
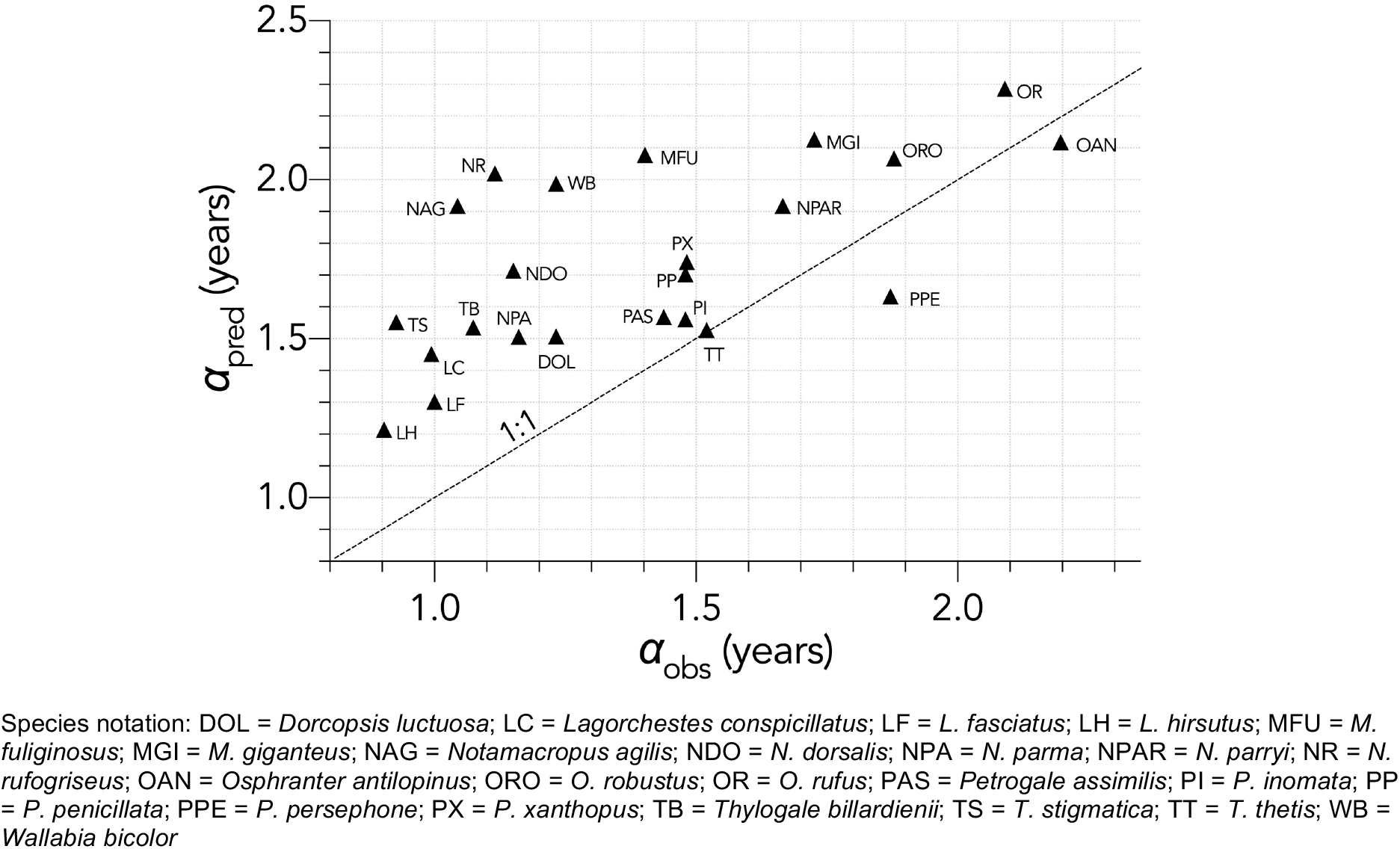
Relationship between the predicted age (years) at first breeding (*α*_pred_) and observed *α* (*α*_obs_) for 23 extant macropodid herbivores [46]. The allometric prediction over-estimated *α* by and average of ~ 20%. Also shown is the 1:1 line (dashed).

Calculating the average disparity between the predicted and observed *α* across species, the allometric prediction over-estimated *α* by 20% for the macropodiform herbivores. We therefore applied this correction factor to the estimated *α* for the extinct macropodiform species (corrected values in Appendix S2, Table S1). The corrected *α* predicted maximum rate of population growth (*r*_m_) approximately following theoretical expectation (see Fig. S6).

**Figure S6.**
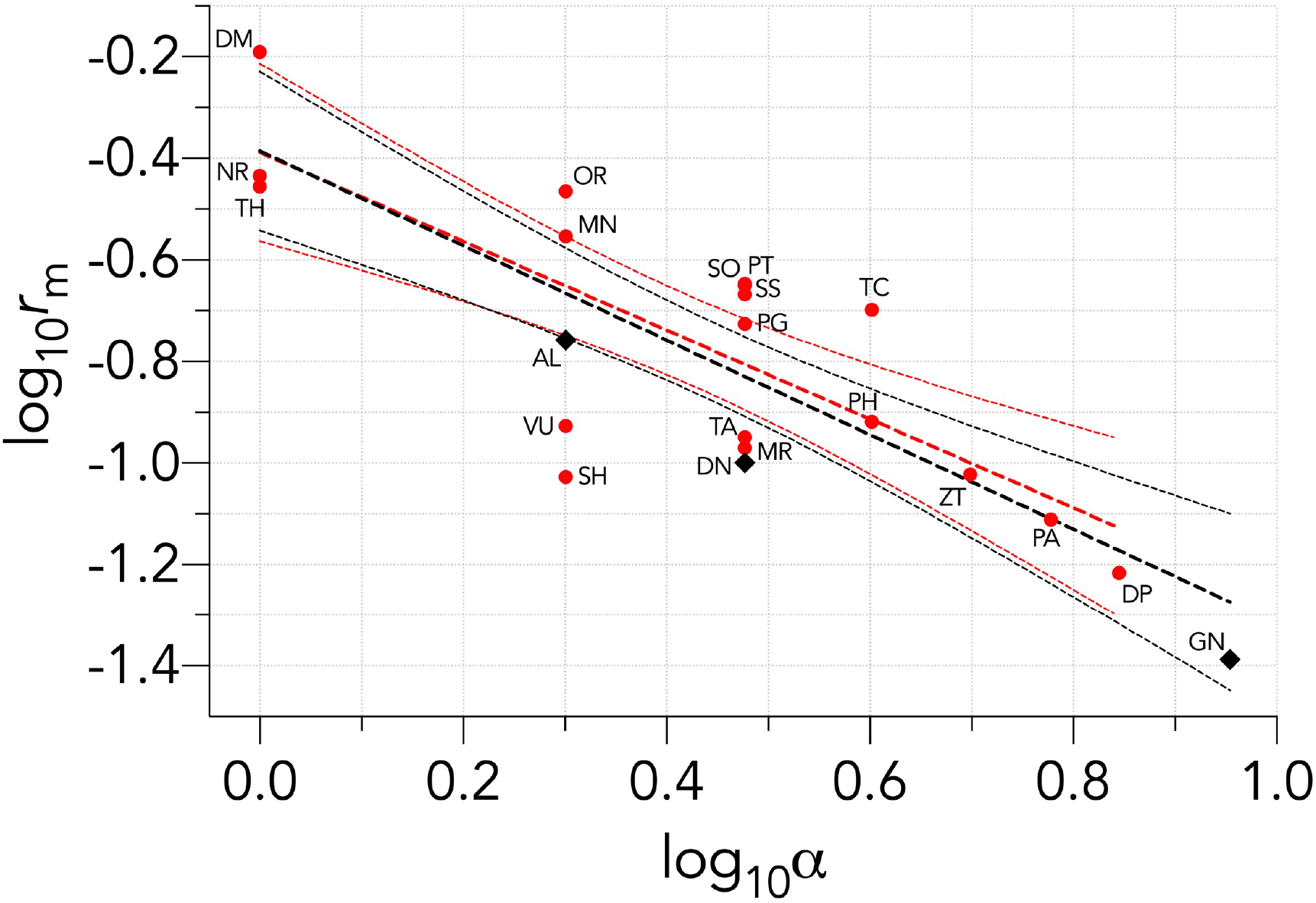
Negative relationship between the logarithm of the maximum rate of intrinsic population growth (log_10_ *r*_m_) and the logarithm of the age at primiparity (log_10_ *α*) for the nineteen species examined. The estimated parameters of the linear fit (*y* ~ *α* + *βx*) including all species (black lines) are: *α* = −0.388 ± 0.075 (± SE) and *β* = −0.931 ± 0.146, and explaining 66.4% of the variation 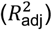, with the information-theoretic evidence ratio (ER) of the slope *versus* intercept-only model = 4.052×10^4^. This relationship is similar to the theoretical expectation for the intercept = −0.15 and slope = −1.0 for mammals[57]. Birds (AL, DN, GN; 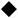) potentially fall outside this relationship, so just considering the remaining mammals 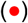, the parameters for the linear fit (red lines) become: *α* = −0.388 ± 0.083 (± SE), *β* = −0.875 ± 0.170, 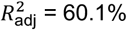, and ER = 1.567×10^3^. Species notation: DP = *Diprotodon optatum*, PA = *Palorchestes azael*, ZT = *Zygomaturus trilobus*, PH = *Phascolonus gigas*, VU *Vombatus ursinus* (**vombatiform herbivores**); PG = *Procoptodon goliah*, SS = *Sthenurus stirlingi*, PT = *Protemnodon anak*, SO = *Simosthenurus occidentalis*, MN = *Metasthenurus newtonae*, OR = *Osphranter rufus*, NR = *Notamacropus rufogriseus*, common name: red-necked wallaby (**macropodiformes**); GN = *Genyornis newtoni*, DN = *Dromaius novaehollandiae*, AL = *Alectura lathami* (**large birds**); TC = *Thylacoleo carnifex*, TH = *Thylacinus cynocephalus*, SH = *Sarcophilus harrisii*, DM = *Dasyurus maculatus* (**carnivores**); TA = *Tachyglossus aculeatus*, MR = *Megalibgwilia ramsayi* (**monotreme invertivores**).

## Appendix S6. Quasi-extinction curves for each species in each of the six perturbation scenarios (Scenarios *ii–vii*)

**Figure S7.**
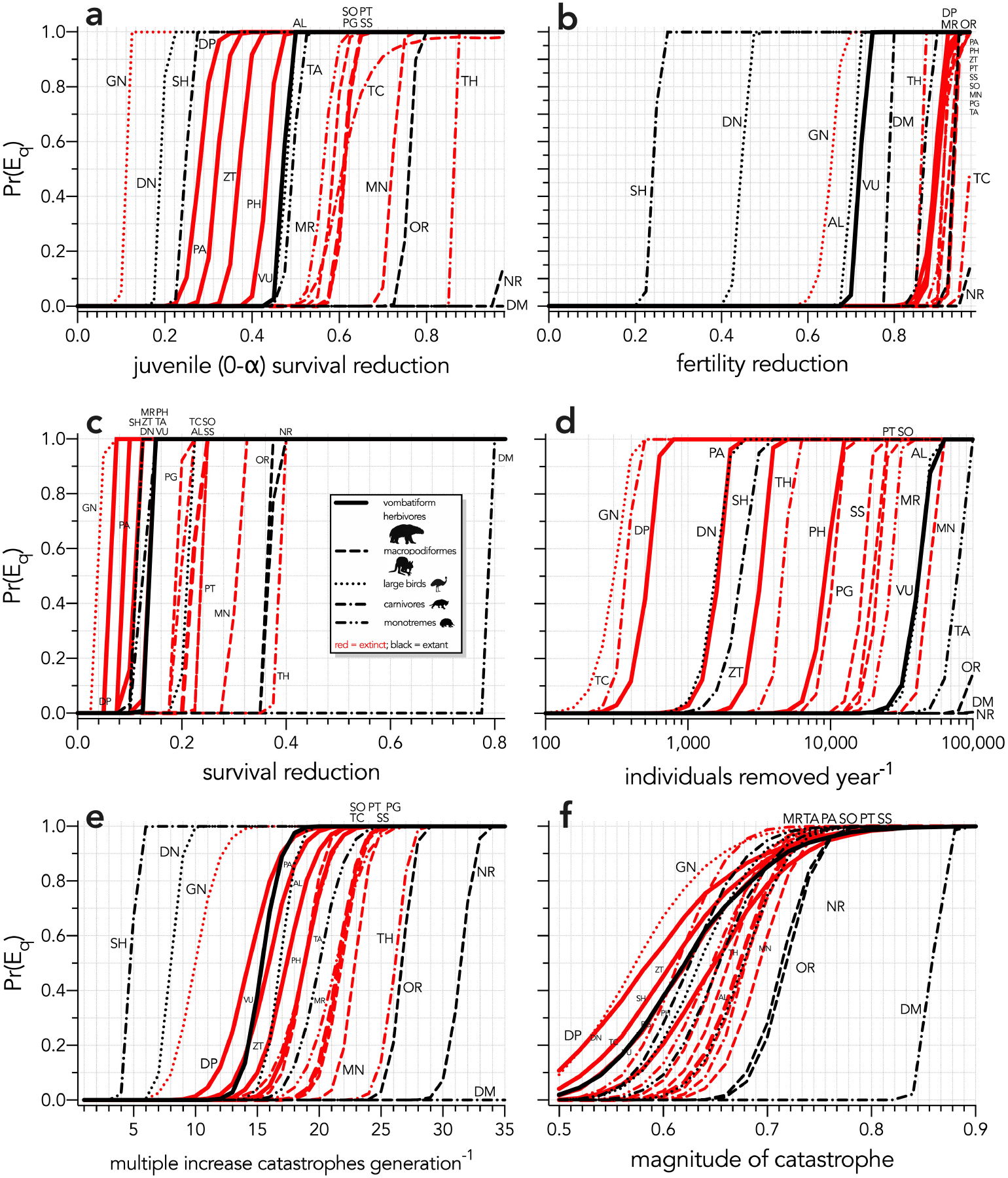
Increasing probabilities of quasi-extinction — Pr(E_q_) — as a function of (a) increasing juvenile mortality (Scenario *ii*), (b) decreasing fertility (Scenario *iii*), (c) decreasing survival across all age classes (Scenario *iv*), (d) increasing number of individuals removed year^-1^ (Scenario *v*), (e) increasing frequency of catastrophic die-offs per generation (Scenario *vi*), and (f) increasing magnitude of catastrophic die-offs (Scenario *vii*). Species notation: DP = *Diprotodon optatum*, PA = *Palorchestes azael*, ZT = *Zygomaturus trilobus*, PH = *Phascolonus gigas*, VU *Vombatus ursinus* (**vombatiform herbivores**); PG = *Procoptodon goliah*, SS = *Sthenurus stirlingi*, PT = *Protemnodon anak*, SO = *Simosthenurus occidentalis*, MN = *Metasthenurus newtonae*, OR = *Osphranter rufus*, NR = *Notamacropus rufogriseus*, common name: red-necked wallaby (**macropodiformes**); GN = *Genyornis newtoni*, DN = *Dromaius novaehollandiae*, AL = *Alectura lathami* (**large birds**); TC = *Thylacoleo carnifex*, TH = *Thylacinus cynocephalus*, SH = *Sarcophilus harrisii*, DM = *Dasyurus maculatus* (**carnivores**); TA = *Tachyglossus aculeatus*, MR = *Megalibgwilia ramsayi* (**monotreme invertivores**). Here, we have depicted SH as ‘extant’, even though it went extinct on the mainland > 3000 years ago.

## Appendix S7. Comparing increasing mortality in juveniles and increasing offtake of juvenile individuals

**Figure S8.**
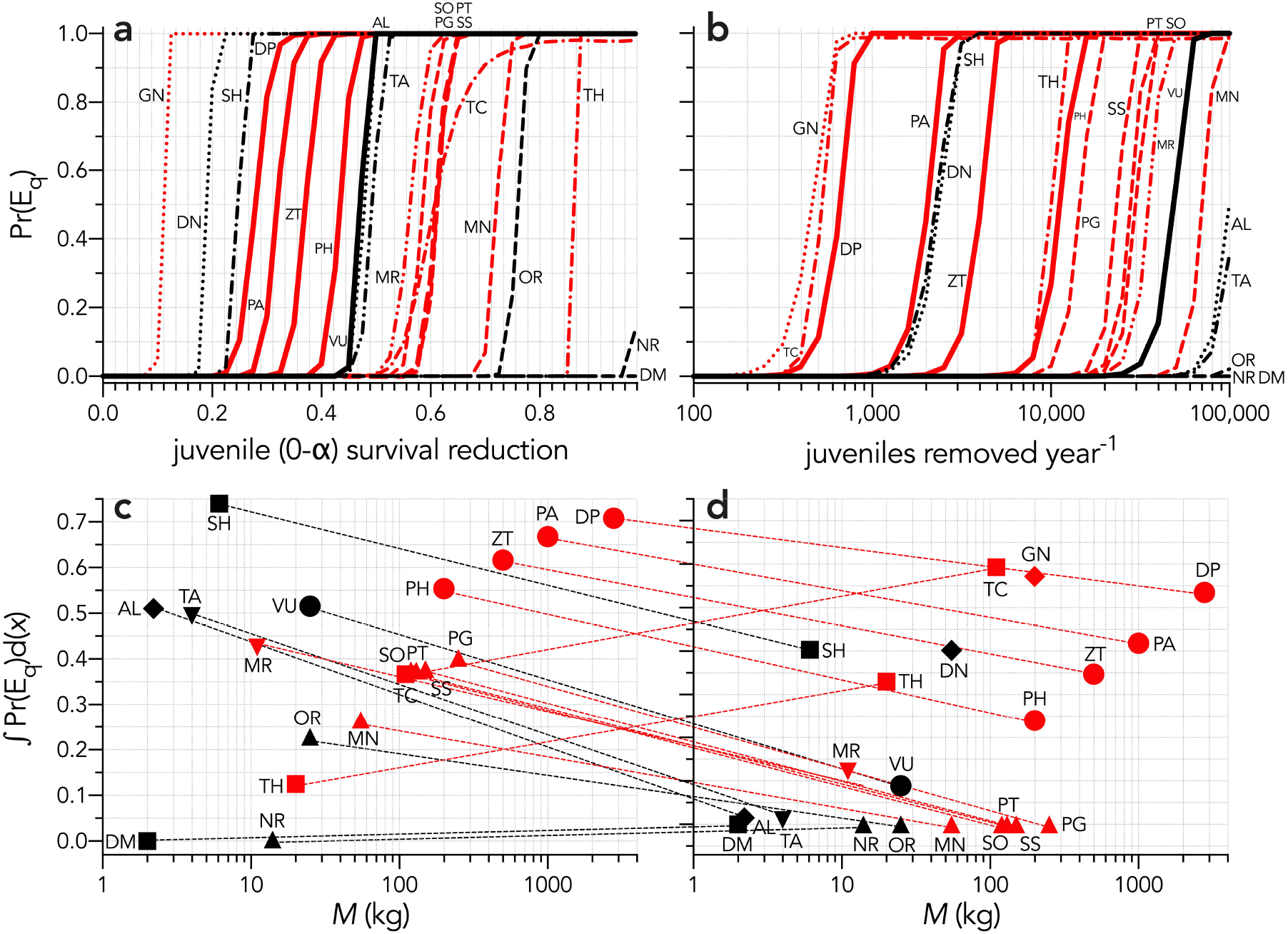
Increasing probabilities of quasi-extinction — Pr(E_q_) — as a function of (a) increasing juvenile mortality (Scenario *ii*), and (b) increasing number of juvenile individuals removed year^-1^ (Scenario *iib*). Also shown is the corresponding area under the quasi-extinction curve — ∫Pr(E_q_)d(*x*) — as a function body mass for (c) increasing juvenile mortality and (d) increasing number of juvenile individuals removed year^-1^. The dashed lines in c and d indicate the change in relative susceptibility between scenarios. Species notation: DP = *Diprotodon optatum*, PA = *Palorchestes azael*, ZT = *Zygomaturus trilobus*, PH = *Phascolonus gigas*, VU *Vombatus ursinus* (**vombatiform herbivores**); PG = *Procoptodon goliah*, SS = *Sthenurus stirlingi*, PT = *Protemnodon anak*, SO = *Simosthenurus occidentalis*, MN = *Metasthenurus newtonae*, OR = *Osphranter rufus*, NR = *Notamacropus rufogriseus*, common name: red-necked wallaby (**macropodiformes**); GN = *Genyornis newtoni*, DN = *Dromaius novaehollandiae*, AL = *Alectura lathami* (**large birds**); TC = *Thylacoleo carnifex*, TH = *Thylacinus cynocephalus*, SH = *Sarcophilus harrisii*, DM = *Dasyurus maculatus* (**carnivores**); TA = *Tachyglossus aculeatus*, MR = *Megalibgwilia ramsayi* (**monotreme invertivores**). Red = extinct; black = extant. Here, we have depicted SH as ‘extant’, even though it went extinct on the mainland > 3000 years ago.

## Appendix S8. Comparing demographic susceptibility to climate variation

Failing to observe any relationship between overall demographic susceptibility and the extinction chronology for Sahul, we might alternatively expect species’ demographic susceptibility to align with increasing environmental stress expressed as hotter, drier climates [66, 67]. We therefore hypothesized that the most extreme (hottest/driest) climates of the past would eventually drive the most-resilient species to extinction, which would manifest as a negative relationship between demographic susceptibility and warming/drying conditions (i.e., only when conditions became bad enough did the least-susceptible species succumb).

To test this hypothesis, we compiled climate indices hindcasted for the estimated extinction windows for the species we considered here. To this end, we acquired four hindcasted, continentally averaged climate variables from the intermediate-complexity, three-dimensional, Earth-system model known as LOVECLIM [68, 69]. LOVECLIM hindcasts various climatic conditions by incorporating representations of the atmosphere, ocean and sea ice, land surface (including a vegetation submodel), ice sheets, and the carbon cycle. These variables were — mean annual temperature (°C), mean annual precipitation (mm), net primary production (kg C m^-2^ year^-1^), and fraction of the landscape designated as ‘desert’ — all expressed as anomalies of their respective average values calculated relative to 120 ka (i.e., a time when all species we considered were extant). We downscaled the original spatial resolution of LOVECLIM (5.625° × 5.625°) to an output resolution of 1° × 1° using bilinear interpolation because it retains the integrity and limitations of the original model output data.

We then calculated the information-theoretic evidence ratios (ER) for all relationships between the mean value of the climate variable across the entirely of Sahul and the sum of the extinction integrals across scenarios as the Akaike’s information criterion (AIC) of the slope model: *y* = *α* + *βx* divided by the AIC of the intercept-only model: *y* = *α* (i.e., ER_mean_ = AIC_slope_/AIC_intercept_). To incorporate full uncertainty in the climate variables (*y*), we developed a randomization test where we uniformly resampled the *y* values between *y*_min_ and *y*_max_, estimating the residual sum of squares of the resampled values at each iteration compared to a randomized order of these residuals. We then calculated the probability (*p*_u_) of producing a randomly generated relationship between the climate variable and the sum of the extinction integrals as the number of iterations when the randomized order produced a residual sum of squares ≤ the residual sum of squares of the resampled (ordered) climate variables divided by the total number of iterations (10,000).

When we plotted the four climate variables against the sum of the quasi-extinction integrals across scenarios, there was evidence for a weak, negative relationship between mean annual precipitation anomaly and relative extinction susceptibility (ER_mean_ = 9.6; Fig. S9b), and a weak, positive relationship with desert-fraction anomaly, (ER_mean_ = 4.2; Fig. S9d). There was no evidence for a relationship between the means for temperature (Fig. S9a) and net primary production (Fig. S9c) anomalies and quasi-extinction integrals. The relationship with desert fraction supports the hypothesis that a drier (more desert-like) environment might have been related to extinction susceptibility. However, when we took full uncertainty of the climate variables into account in a randomization leastsquares regression, none of the relationships could not be differentiated from a random process (*p*_u_ > 0.17; Fig. S9).

**Figure S9.**
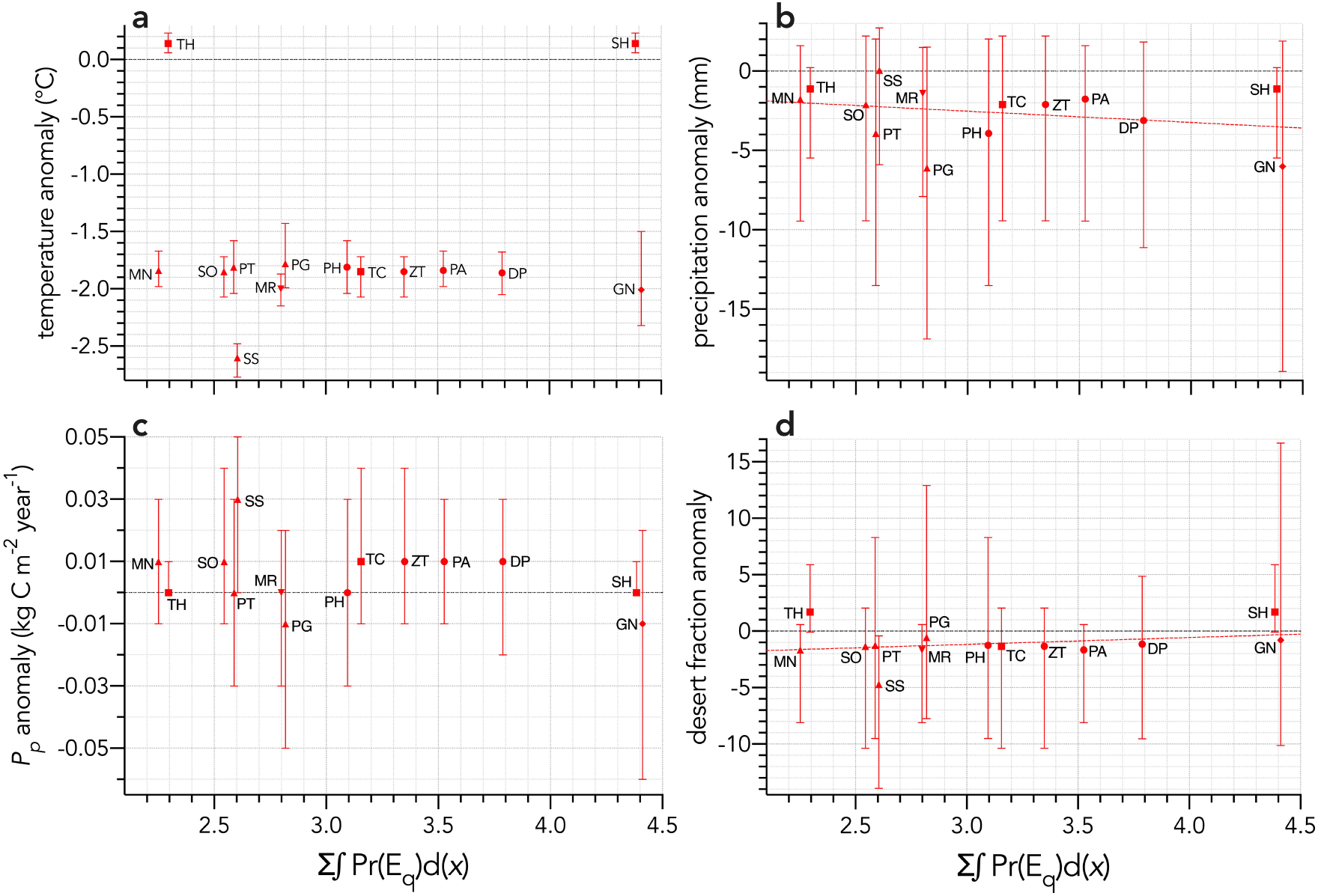
Sum of the areas under the quasi-extinction curve over the six scenarios considered — Σ∫Pr(E_q_)d(*x*) — for each of the 13 extinct (mainland only) modelled species relative to (**a**) mean annual temperature anomaly (°C): information-theoretic evidence ratio of the slope model relative to the intercept-only model (ER_mean_) = 0.75 for the mean climate values; probability of a non-random slope relationship incorporating full uncertainty in the climate variable *p*_u_ = 0.603. (**b**) mean annual precipitation anomaly (mm): ER_mean_ = 9.6; *p*_u_ = 0.461. (**c**) net primary production anomaly (kg C m^-2^ year^-1^): ER_mean_ < 0.01; *p*_u_ = 0.411. (**d**) desert fraction anomaly: ER_mean_ = 4.2; *p*_u_ = 0.425. The dashed red line in panels a, b, and d indicate evidence for a slope model versus the intercept-only model for these variables (ER_mean_ > 2). Error bars indicate ± 1 standard deviation. Species notation: DP = *Diprotodon optatum*, PA = *Palorchestes azael*, ZT = *Zygomaturus trilobus*, PH = *Phascolonus gigas* (**vombatiform herbivores**); PG = *Procoptodon goliah*, SS = *Sthenurus stirlingi*, PT = *Protemnodon anak*, SO = *Simosthenurus occidentalis*, MN = *Metasthenurus newtonae* (**macropodiformes**); GN = *Genyornis newtoni* (**large bird**); TC = *Thylacoleo carnifex*, TH = *Thylacinus cynocephalus*, SH = *Sarcophilus harrisii* (**carnivores**); MR = *Megalibgwilia ramsayi* (**monotreme**).

The lack of evidence for a relationship between extinction susceptibility and warming/drying conditions for the mean climate conditions across the continent contrasts with recent evidence that water availability potentially exacerbated mortality from novel human hunting [66, 67]. However, there is too much uncertainty in the climate hindcasts to test this hypothesis definitively. Another weakness of this approach is that we were obliged to take continental-scale averages of average climate conditions, which obviously ignores spatial complexity previously established as an important element in explaining the chronology and directionality of megafauna extinctions, at least in south-eastern Sahul [66, 67]. Thus, this enticing, but still unsupported hypothesis that warming and drying conditions were related to intrinsic extinction susceptibility, will require more precise estimates of extinction timing and hindcasted climate conditions, and perhaps greater sample sizes across more species.

